# *Clostridial* acidogenesis product profiles are discontinuous: a thermodynamic hypothesis

**DOI:** 10.64898/2026.06.03.729792

**Authors:** Christoff Odendaal, Marit Verheijen, Rebeca Gonzalez-Cabaleiro

## Abstract

At neutral pH, *Clostridial* fermentative catabolism is typically acidogenic, with a product profile dominated by acetate and butyrate. H_2_ acts as a terminal electron acceptor via hydrogenases, which increases ATP-producing potential from glucose. Acetate production is characterised by both higher ATP and H_2_ yields, rendering it desirable but more thermodynamically limited. For this reason, is commonly understood that *Clostridia* can adjust the ratio of acetate to butyrate (Ace:But) produced to maximise ATP while maintaining sufficient pathway driving forces to sustain a high flux.

We identify three redox-balanced product profiles that underlie the spectrum of *Clostridial* catabolic Ace:But ratios: Homoacetic (Ace:But = 2:0), Equimolar (0.67:0.67), and Homobutyric (0:1). To reach Ace:But ratios intermediate to these, the elementary flux modes (EFMs) underlying the aforementioned product profiles must be blended.

We performed a maximum-minimum driving force (MDF) analysis to test the thermodynamic favourability of the pathways underlying different Ace:But ratios at varying H_2_ partial pressures (pH_2_). We find that blended EFMs are less efficient than their constituent EFMs at all pH_2_, allocating excessive driving force (DF) to certain reactions, thereby lowering the DF of others. This is, in part, due to the co-occurrence of hydrogenases with different optimal redox carrier ratios. One hydrogenase inevitably has very high DF, which decreases the DF available for other reactions. This leads to a lower minimum DF and a higher enzyme cost for operating blended EFMs. This implies that certain discrete Ace:But ratios are most favourable for large ranges of pH_2_, contradicting the continuity assumption in literature.

## Introduction

Anaerobic fermentations are metabolic processes where organic compounds are broken down in the absence of an external electron acceptor. Consequently, the substrate serves as both electron donor and acceptor to maintain the metabolic redox balance. Glucose fermentation, for example, begins with glycolysis, in which the substrate is oxidised to pyruvate. To provide additional growth precursors, pyruvate is decarboxylated and oxidised to acetyl-CoA, concomitantly yielding either CO_2_ and H_2_, or formate. The requirement to maintain internal redox balance constrains the number of feasible catabolic routes in fermentative glucose metabolism. In these pathways, reducing equivalents are primarily transferred via two key electron carriers: nicotinamide adenine dinucleotide, NADH (-320 mV), and ferredoxin, Fd(red) (-400 mV)^1^. Depending on the route through which these carriers are reoxidised, various volatile fatty acids (VFAs) and alcohols can be formed. Acetate and butyrate are dominant end products commonly observed in anaerobic glucose-fermenting systems dominated by *Clostridium* species ^2– 6^.

During such fermentations, ferredoxin is reoxidised via proton reduction to hydrogen gas (H_2_) as the terminal electron acceptor. This happens via a collection of four redox-balancing reactions. The first of these entail hydrogenases that simply produce H_2_ from Fd(red), variously referred in the literature as “non-bifurcating hydrogenases” ^7^, “[FeFe]-hydrogenases” ^8^, “clostridial-type [FeFe] hydrogenases” ^9,10^, and “monomeric [FeFe]-hydrogenases of Group A” ^9^. For simplicity, we will refer to them as “hydrogenase (Hyd)”. Secondly, there are “confurcating hydrogenases” (Conf_Hyd) that confurcate electrons from Fd(red) and NADH to produce H_2_. Thirdly, the “energy-converting hydrogenases” (EC_Hyd) couple hydrogen production to the translocation of one proton (or Na^+^) out of the cytosol into the periplasm, where it can be used for ATP production via the ATP synthase ^11,12^. It is estimated that this yields an additional 0.25 molecules of ATP per reaction, equivalent to 1 proton pumped per electron transferred ^8,11^. Fourthly, the RNF-complex (ferredoxin:NAD+ oxidoreductase) couple the oxidation of Fd(red) to the reduction of NAD^+^; the Gibbs free energy change of this oxidation also allows for a transmembrane ion gradient (Δ µH+/Na+) that can drive additional ATP-synthesis ^13,14^.

Various *Clostridia* have been shown to contain and express these four redox-balancing enzymes in different combinations ^13,15–21^. The presence of these different enzyme bundles would allow species of this class to flexibly use different reductive routes in their catabolism, ranging from homoacetic to homobutyric fermentation and any linear combination between those two extremes. ATP can be produced either via substrate-level phosphorylation (SLP) or transport-coupled phosphorylation (TCP), the latter depending on the formation of ion-motive force, usually proton-motive force (PMF) ^4^. Since PMF can be dissipated or used for other purposes ^22,23^, ATP production from SLP can be regarded as the only numerically certain ATP yield from catabolism. Acetate is associated with increased ATP yield via SLP (4 ATP/Gluc), but also increased H_2_ production, while butyrate yields lower ATP (3 ATP/Gluc) but also less H_2_ (**Fig. 1**).

**Figure 1.**
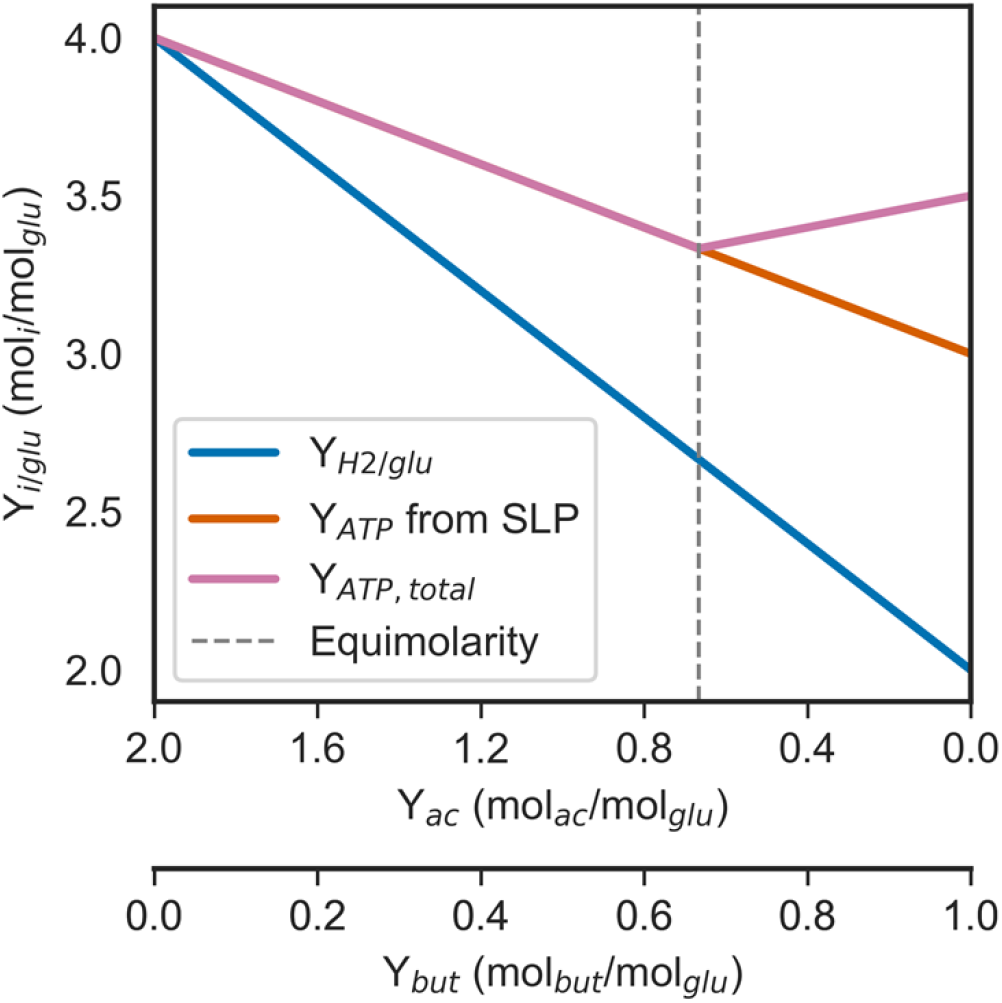
H_2_ production and ATP yield as a function of the fermentation product mix. The product ratio between acetate and butyrate is depicted on the x-axis, with H_2_ and ATP production on the y-axis. ATP production is highlighted as the total production (pink) and the contribution from substrate-level phosphorylation (SLP) alone. The total production includes ATP resulting from transport-coupled phosphorylation, assuming a ratio of 4 H^+^/ATP for ATP synthase and that the RNF-complex translocates two ions per turnover. Equimolar production of acetate and butyrate is indicated by the dashed vertical line.

It is frequently assumed that the ratio of acetate:butyrate produced from glucose (Ace:But ratio) flexibly adapts to the hydrogen partial pressure (pH_2_) to maximise ATP production and that the “transitions between [pure acetate and pure butyrate production] are continuous”, with acetate dominating the product profile at low pH_2_ and butyrate at high pH_2_ ^4^. However, this is not reflected in experimental Ace:But ratios from either mixed or pure *Clostridial* cultures from literature where the pH_2_ is varied (**Fig. S1**): the Ace:But ratio, especially of fast-growing cultures – tends to be relatively insensitive to changes in pH_2_ ^24–27^. This aligns with the fact that the literature contains examples of homoacetic (in environments with pH_2_ ≤10 Pa) ^28^, and of equimolar glucose-catabolic stoichiometries ^1,7,13^, but not of combinations. This is remarkable, considering the observed interplay between pH_2_ sensitivity and ATP yield (**Fig. 1**).

To explore the feasibility of these different glucose-catabolic stoichiometries, we applied a thermodynamic framework to examine the driving forces underlying the various stoichiometries ranging between homoacetic and homobutyric fermentation. *Noor et al*. ^29^ have described the Flux-Force relationship or Flux-Force Efficacy (FFE), highlighting increasing back flux at decreasing driving force, which translates to higher enzyme cost. Stoichiometries that display higher FFE for most reactions incur a lower protein cost. The relationship between FFE and DF is hyperbolic, and DF ≥10 kJ/mol has been shown to only negligibly increase FFE ^29^. This DF is then unavailable to other reactions, so we refer to it as “wasted DF”.

We argue that the ease of distributing DF is key in determining the favourability of a fermentative pathway stoichiometry. As a key example, we show that combining the homoacetic and equimolar catabolic stoichiometries to increase ATP yield per glucose leads to DF distributions that can give rise to increased enzyme cost per ATP produced.

## Results

### Catabolic stoichiometries in *Clostridia*

First, we constructed a network including the central acetate-butyrate-producing catabolic reactions present in *Clostridia* (**Table S3**) ^4,9^. We then identified the redox-balanced stoichiometries within the network using elementary flux mode (EFM) enumeration ^30^. EFMs are the simplest balanced stoichiometries in a network, so that all balanced stoichiometries must either be EFMs or combinations of EFMs ^31^. We identified three product profiles, for which we display the following representative catabolic stoichiometries (EFMs) in **Figure 2**, with the following overall stoichiometric reactions:

**Figure 2.**
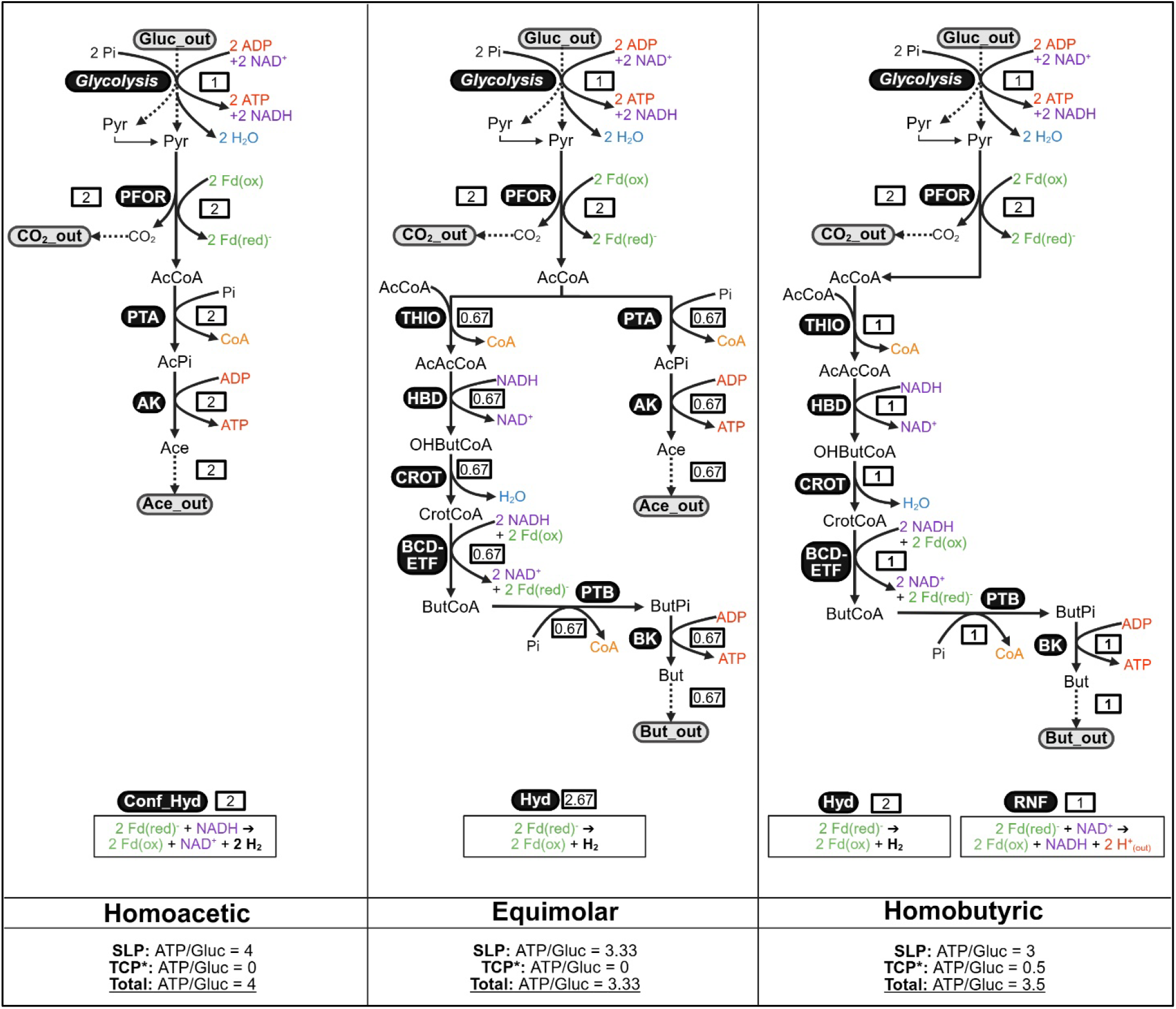
Balanced stoichiometries in *Clostridial* acidogenesis. To identify the balanced stoichiometries in *Clostridial* acidogenesis, a network of catabolic reactions in *Clostridia* was decomposed into elementary flux modes (EFMs). Three product profiles were identified with different Ace:But ratios: Homoacetic (2:0), Equimolar (0.67:0.67), and Homobutyric (0:1). The figure displays a representative EFM for each of these conversions. Black, curved boxes contain enzyme name abbreviations as defined in the glossary and grey curved boxes, the extracellular metabolites. ATP and pumped protons (red), redox equivalents (NADH/ NAD^+^ in purple and Fd(red)/Fd(ox) in green), the conserved moiety free CoA (orange), and H_2_O (fixed concentration, blue) are coloured in, for discernibility. H_2_, the final electron acceptor from the Fd(red), is in bold text. ATP production is displayed at the bottom of each EFM: ATP from substrate-level phosphorylation (SLP), from transport-coupled phosphorylation (TCP). The asterisk (*) next to TCP indicates that it is an upper limit: we assume full utilisation of proton motive force (PMF) for ATP production. Dashed arrows indicate condensed steps of glycolysis and transport. Alternative EFMs were identified for Equimolar and Homobutyric, with variations in ATP yield and redox-balancing enzymes used. These are shown in **Figure S2**. The full stoichiometries, expanded with the glycolysis reactions, are in **Table S2**.

**Homoacetic:**

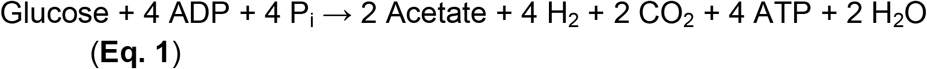

**Equimolar:**

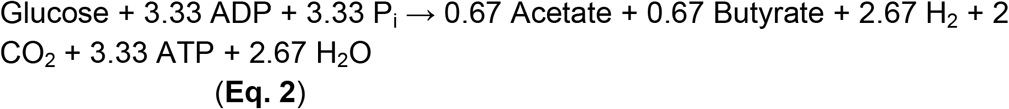

**Homobutyric:**

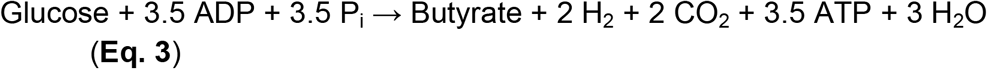

ATP consumption was attributed exclusively to non-catabolic processes, hence the stoichiometries in **Fig. 2** present it as a product. ATP can be conserved either via substrate-level phosphorylation (SLP) or transport-coupled phosphorylation (TCP). The former decreases as a function of the H_2_ production of the pathway. TCP – which generates ATP by first generating proton-motive force (PMF) – does not necessarily follow this trend (**Fig. 1**). However, PMF is a less directly quantifiable source of ATP because it can also drive other cellular processes (e.g. proton-coupled symport ^22^ or dissipation through proton leakage ^23^). Therefore, TCP and SLP are presented separately before being summed in **Fig. 2**.

### Analysis of driving force distributions

A Max-Min Driving Force (MDF) analysis identifies the combination of metabolite concentrations within a pathway which maximises the lowest reaction driving force (DF) across all reactions ^29^. **Fig. 3** displays the full result of the MDF analysis for one of the identified EFMs, an equimolar acetic/butyric stoichiometry (**Fig. 3A**) ^2^. **Fig. 3B** contains the metabolite concentrations returned by the MDF analysis in molar (M) units. The MDF non-linear optimisation is not unique. Error bars represent the calculated variability in the metabolite concentrations for which the MDF objective solution is valid.

**Figure 3.**
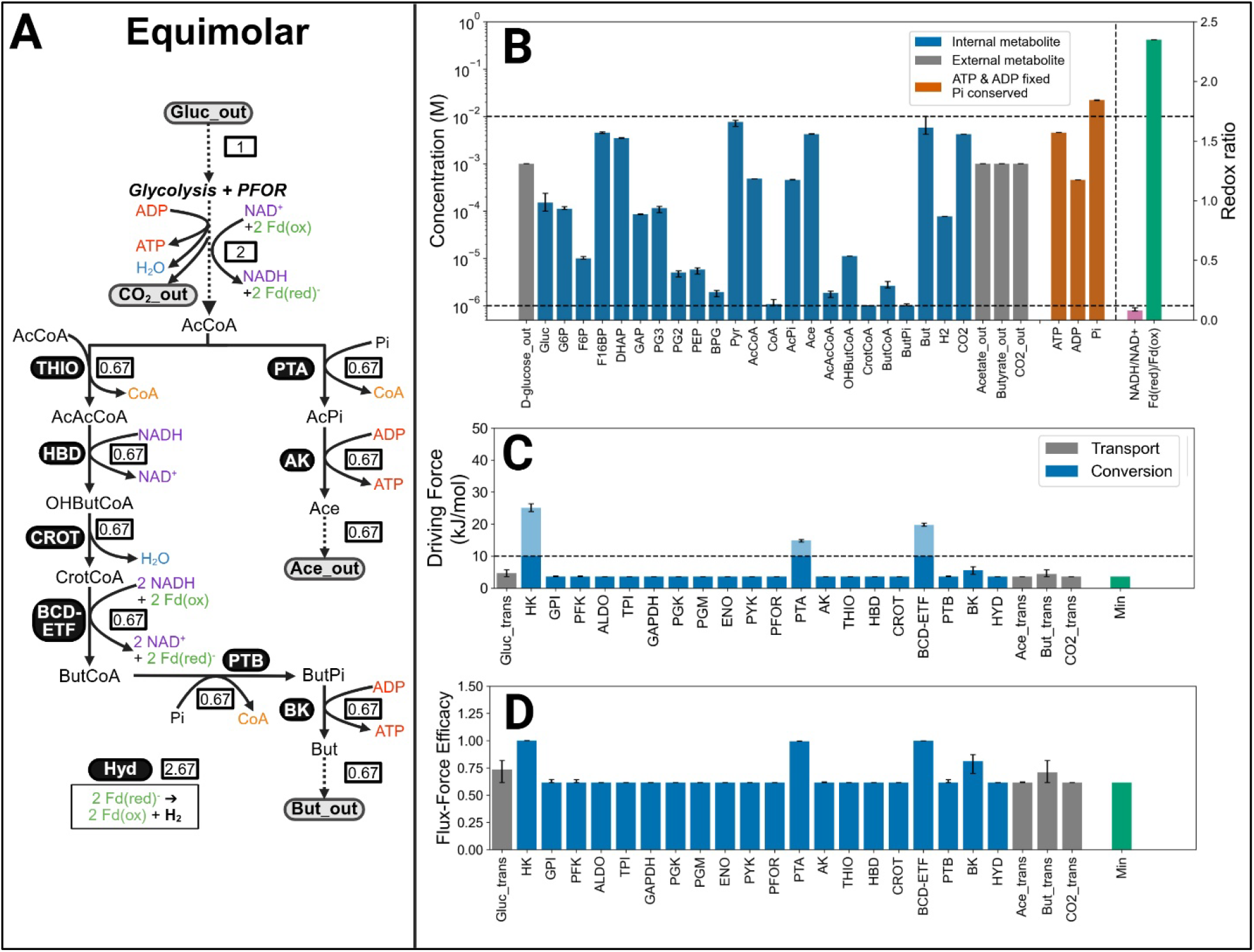
Max-Min Driving Force analysis of Equimolar. An illustrative example of a Max-Min Driving Force (MDF) analysis: detailed results pertaining to the product profile associated with *Clostridium pasteurianum* at pH_2_ = 10 kPa. The error bars represent the ranges for which the minimum DF remains optimal and the horizontal dashed lines show the concentration bounds or all variable metabolites except P_i_ and CoA (see *Methods*). **A**. Flux distribution of Equimolar. **B**. Metabolite concentrations (left-hand y-axis) and redox ratios (right-hand y-axis). External metabolites and ATP/ADP have fixed concentrations. Total Pi, like total CoA, is a conserved moiety. NADH/NAD^+^ and Fd(red)/Fd(ox) are displayed as ratios. **C**. Driving forces (DF) allocated to each reaction. DF > 10 kJ/mol has negligible effect on flux force efficacy (FFE) and are presented as faded. Transport and conversion reactions are distinguished. The green bar shows the maximised minimum DF. **D**. FFE calculated from the driving forces in panel **C**: 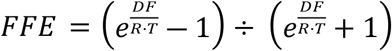.

**Figure 3C** presents the reaction driving force (DFs, in kJ/mol) distribution obtained from maximising the pathway minimum DF. For this catabolic stoichiometry, the DF distribution corresponds to a flux-force efficacy (FFE) all well above 60% for all the reactions of the pathway (**Fig. 3D**). The comparison of DF and FFE values (**Fig. 3C-D**) shows how a large difference in DF (e.g. between HK and GPI – a 4-fold difference) translates to only an approximately 30% difference in FFE, as FFE increases asymptotically as a function of DF. Beyond DF > 10 kJ/mol, additional DF provides negligible changes in FFE ^29^. We therefore interpret DFs above this threshold as excess thermodynamic investment (energy dissipated that do not contribute to higher metabolic fluxes) and use them to quantify pathway inefficiency. Throughout this paper, this is referred to as ‘Wasted DF’.

### DF distributions and FFE as a function of pH_2_

To understand how the EFMs became more thermodynamically favourable at different H_2_ partial pressures (pH_2_), we performed MDF analyses on the three representative catabolic stoichiometries at pH_2_ ranging from 10 Pa to 1000 kPa. **Fig. 4A** shows the distribution of DF across reactions at pH_2_ = 10 kPa. Note that the stoichiometries have different lengths: Homoacetic consists of 18 reactions (including transport), Equimolar of 24, and Homobutyric of 20. **Fig. 4B** shows how the different stoichiometries have different total pathway driving forces at various pH_2_. As all these stoichiometries are dependent on H_2_ production to reoxidise ferredoxin, they all have decreasing total pathway DF as a function of increasing pH_2_ with a steeper decline for stoichiometries that yield more H_2_.

**Figure 4.**
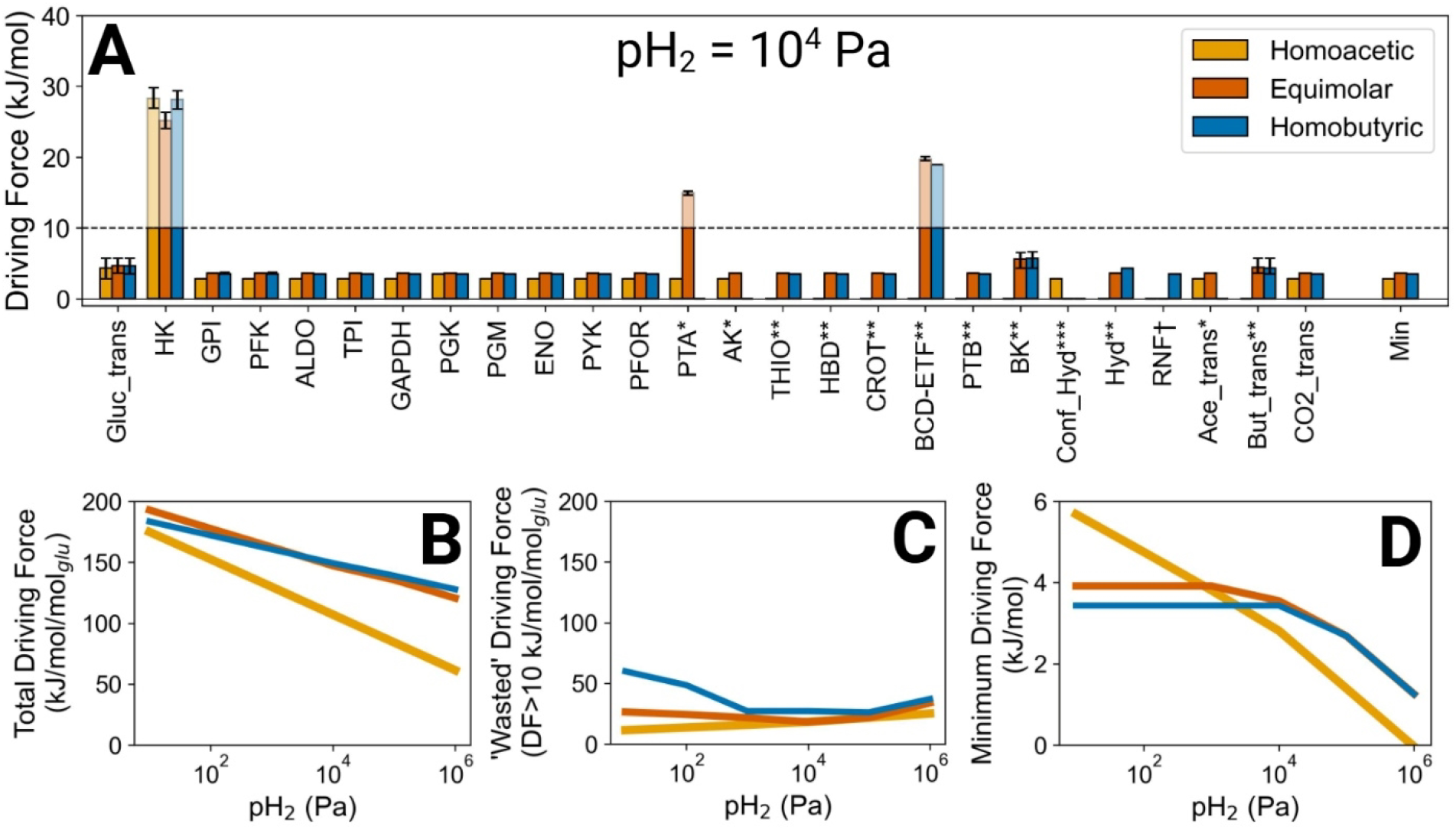
Driving force as a function of pH_2_ in different EFMs. The results of MDF analyses performed on representative *Clostridial* acidogenic catabolic stoichiometries. **A**. Comparison of driving forces per reaction at pH_2_ = 10 kPa. * reactions not in Homobutyric; ** reactions not in Homoacetic; *** reactions only in Homoacetic; † reaction only in Homobutyric. **B**. Total, flux-weighted pathway driving force (DF) in units of kJ/mol/mol_glu_. The DF is the remaining -ΔG_r_’ after the energy of ATP production and proton pumping has been subtracted. **B**. ‘Wasted’ DF – sum of the fraction of DF in each reaction > 10 kJ/mol /mol_glu_. **C**. Minimum DF (kJ/mol) of any reaction stoichiometry, the thermodynamic bottleneck of the pathway.

In contrast to the regularity with which the total pathway DF declines as a function of increasing pH_2_, there is an associated complexity to how these different EFMs distribute their driving force across individual reactions. We defined all DF > 10 kJ/mol as ‘Wasted’ and calculated the flux-weighted sum of ‘Wasted’ DF (**Fig. 4C**). The maximum ‘waste’ is observed at different pH_2_ values for the different EFMs, with Homobutyric being the most inefficient stoichiometry.

Homoacetic has the highest minimum DF at low pH_2_ (**Fig. 4D**), despite having lower total DF. It also has the highest SLP yield, at 4 ATP/Gluc, and the highest H_2_ yield (4 moles of H_2_ per mole of glucose). When pH_2_ increases, the DF available decreases sharply, which reduces DF of the enzymatic reactions, having the lowest minimum DF (<3 kJ/mol at pH_2_ < 10 kPa). This limits the possible flux of the pathway. Although Homoacetic is exergonic at all ranges of pH_2_ (**Fig. 4B**), this constrained minimum DF (**Fig. 4D**) can explain why cultivated homoacetogens are typically extremophiles, growing at pH_2_ < 100 kPa ^8,32^. Equimolar has the highest minimum DF at pH_2_ = 1 – 100 kPa).

Homobutyric has high ‘wasted’ DF and therefore the lowest minimum DF of all product profiles. Nevertheless, at higher values of pH_2_, the minimum DF of Homobutyric reaches similar values to the ones render for Equimolar stoichiometry, as its waste decreases and the total DF becomes the highest of all stoichiometries (**Fig. 4B**).

Note that we also tested the alternative Homobutyric and Equimolar stoichiometries (**Fig. S3**). For they had lower minimum DF than the stoichiometries in the main text (**Fig. S3**), indicating that they are less favourable. For simplicity, we do not go into their MDF results in further detail.

### Driving forces of blended stoichiometries

Evaluating the EFMs investigated in **Fig. 4** showed that the thermodynamic favourability changes continuously as a function of pH_2_. Hypothetically, blends of EFMs might have intermediate ATP yield and thermodynamic favourability at intermediate pH_2_. We investigate two examples:

#### 50% Homoacetic (Eq. 1) + 50% Equimolar (Eq. 2)

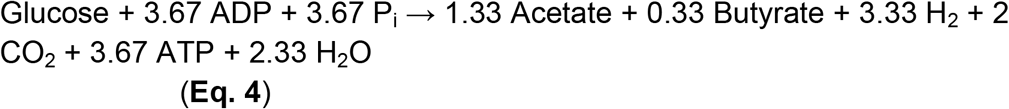

#### 50% Homobutyric (Eq. 3) + 50% Equimolar (Eq. 5)

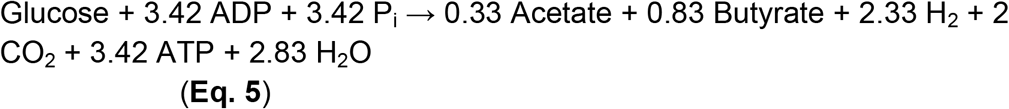

**Fig. 5A-C** shows that while the total DF of the blended stoichiometries was intermediate in both cases (**Fig. 5A**), the “wasted” and minimum DFs were not. The combination Homoacetic+Equimolar (Eq. 4) showed the highest wasted DF values above 1 kPa (**Fig. 5B**) and a minimum DF lower than both of its constituent EFMs at all pH_2_ values (**Fig. 5C**).

**Figure 5.**
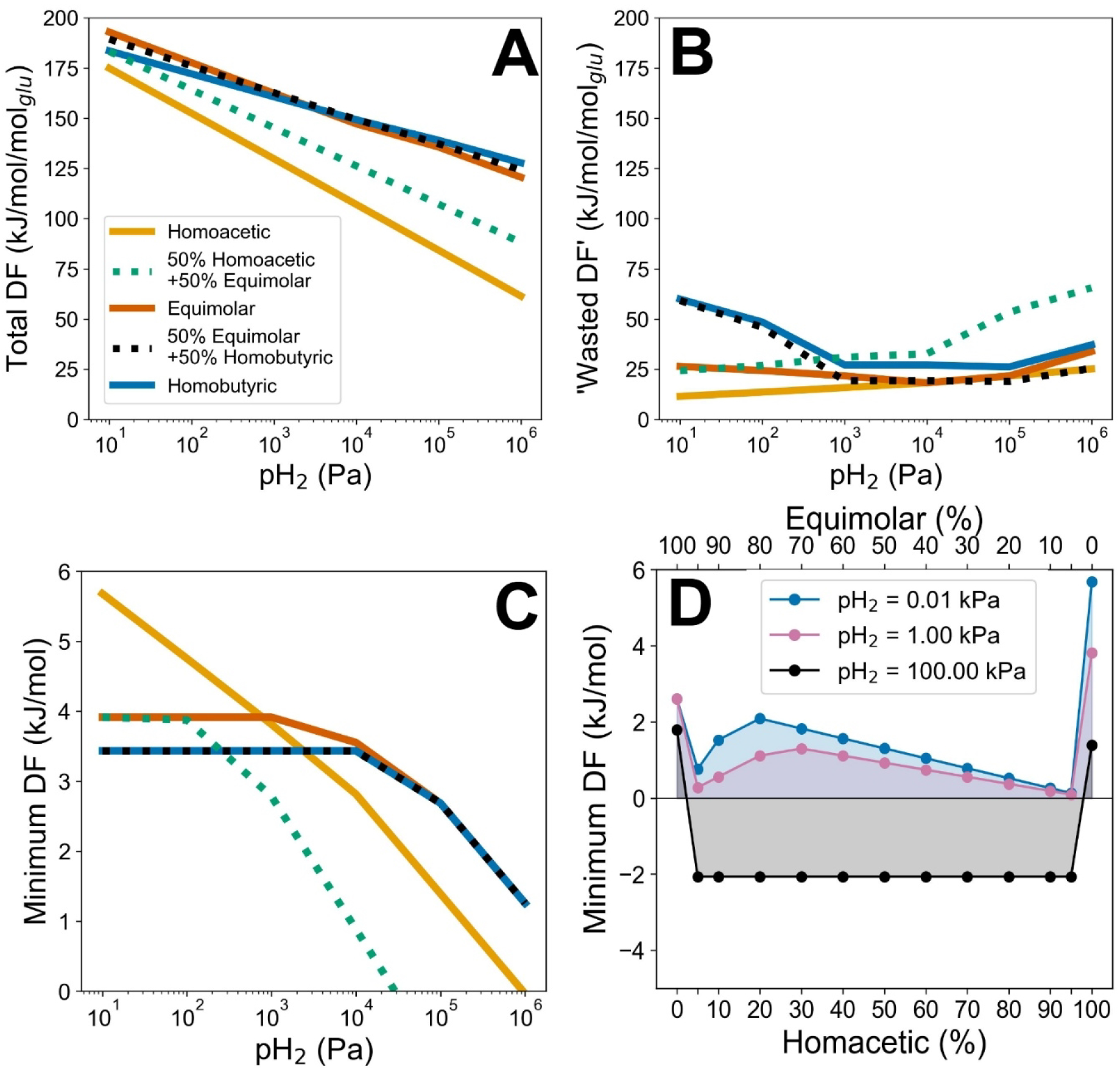
Driving forces of blended stoichiometries. Homoacetic and Equimolar, and Homobutyric and Equimolar were blended in a 50:50 ratio and MDF analyses performed at different pH_2_ values. The total (**A**), ‘wasted’ (**B**), and minimum (**C**) DF are shown for the original and blended stoichiometries. MDF analyses were also performed on Homoacetic and Equimolar blend in other ratios (**D**). Note the two x-axes representing the % of Equimolar (top) and Homoacetic (bottom). MDF was carried out at three different pH_2_ values for all combinations.

Indeed, 15 out of 26 pathway reactions had lower DF in the blended stoichiometry than in either contributing EFM (**Fig. S4**), and 14 of those reactions had a flux-force efficacies (FFEs) that were ≥2-fold lower than in the EFMs. This imposes a substantial enzyme cost on the blended stoichiometry to sustain a similar flux to either independent EFM. Blending Homobutyric and Equimolar did not substantially increase either the “wasted” or minimum DF, but it also did not increased the minimum DF relative to Homobutyric (**Fig. 5C**).

Another possibility is that the blended stoichiometries are thermodynamically superior at certain specific ratios. Since the blend 50% Homoacetic + 50% Equimolar had such a large impact on DF distribution, we extended the analysis to other ratios of combinations of Homoacetic and Equimolar at various ratios; we also performed the analysis at three different pH_2_ values (**Fig. 5D**). Strikingly, even a 5:95 combination in either direction of these two EFMs led to a lower minimum DF than either of the pure EFMs in all cases examined. This coincides with an increased ‘waste’ at all blend ratios (**Fig. S5**), suggesting that any fractional combination of Homoacetic and Equimolar leads to inefficiencies in distributing DF.

### Box 1: It starts with the hydrogenases

The reason why a blend of two EFMs is less favourable than either one of the constituent EFMs (e.g., Homoacetic and Equimolar) starts with the hydrogenases. Homoacetic makes use of a confurcating hydrogenase (Conf_Hyd) which simultaneously reoxidises one NADH and two Fd(red) while producing 4 molecules of H_2_ (**Fig. 2**). This corresponds to the tri- and tertrameric enzymes HydABC ^8^ and HndABCD ^33^:

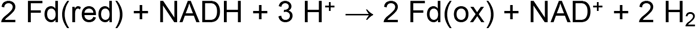

Equimolar, on the other hand, makes use of a hydrogenase which reoxidises two Fd(red) while producing 2.66 molecules of H_2_. This reaction can be carried out by multiple monomeric iron-only hydrogenases (Hyd) ^9^:

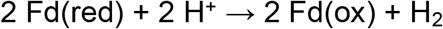

These hydrogenases share most metabolites: Fd(red), Fd(ox), and H_2_. If Conf_Hyd and Hyd co-occur in the same compartment, they also encounter the same concentrations of these shared metabolites, then their flux weighted DFs would be a function of the same Fd(red)/Fd(ox) ratio at a given pH_2_. The DF of Conf_Hyd would be able to vary independently of Hyd’s to a limited degree due to the range of possible values of the ratio NADH/NAD^+^ (typically constrained to a range of 0.01-1 ^8,34–36^):

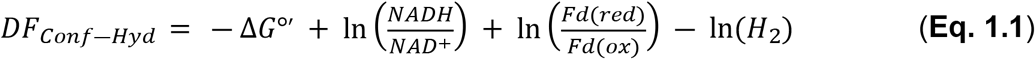

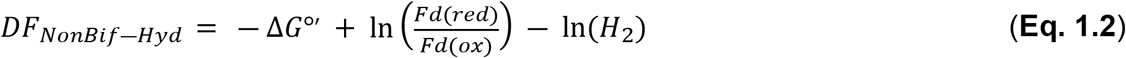

**Fig. 6A** shows this relationship for the physiologically relevant range of Fd(red)/Fd(ox) = 1 - 100 ^8,37^. It shows that Conf_Hyd can reach a positive DF only if Hyd has a flux-weighted DF > 20 kJ/mol. **Fig. 6B** shows that the FFE of Hyd has, by this point, long been saturated, essentially meaning that all additional DF allocated to it is ‘wasted’. This also holds at very high or very low pH_2_ for the same combination (**Fig. S6**) as well as for the combination of Homobutyric and Equimolar at high and low pH_2_ (**Fig. S7**).

**Figure 6.**
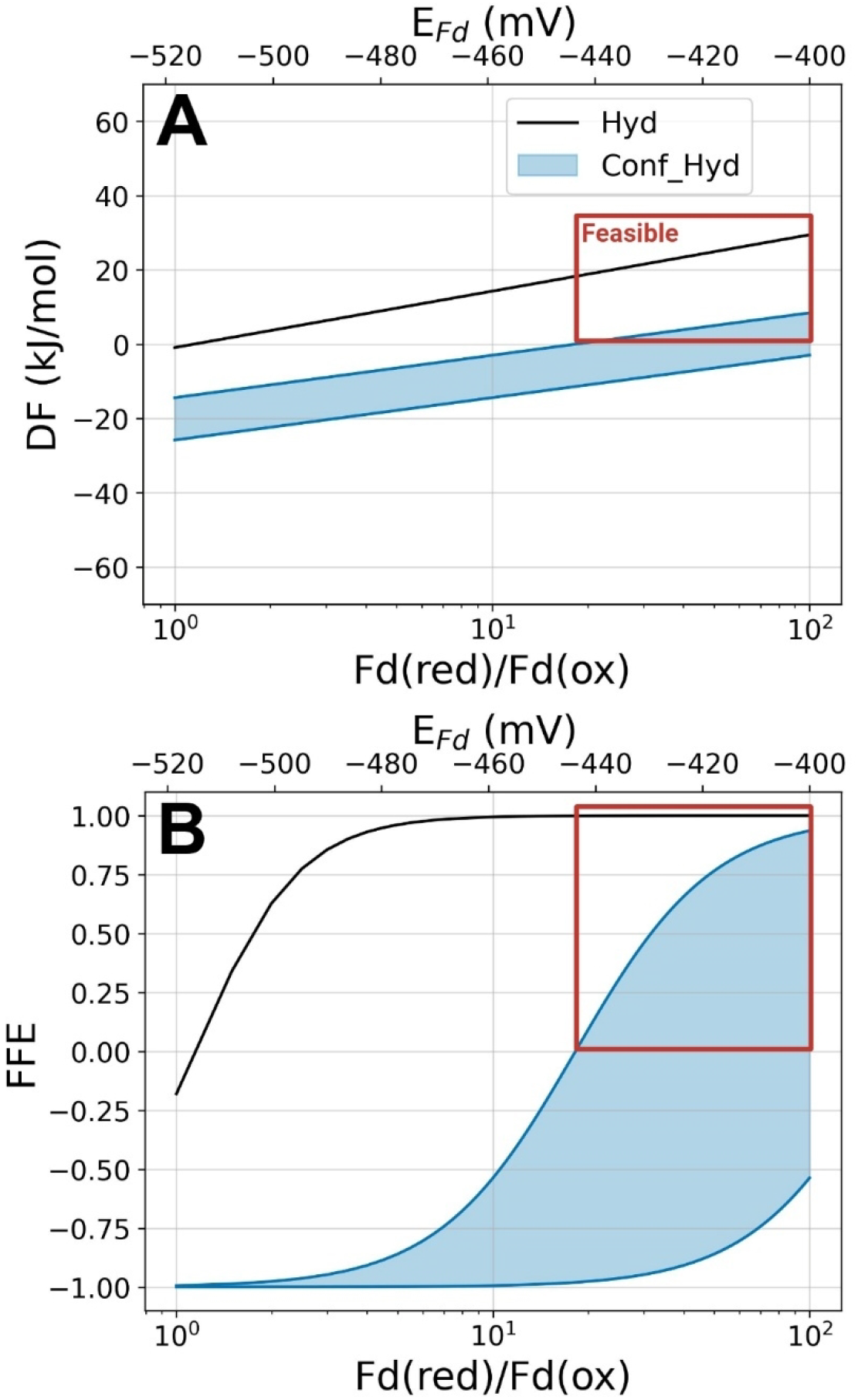
Combining Hyd and Conf_Hyd introduces thermodynamic limitations. The disfavourability of blending Homoacetic and Equimolar stems from the fact that these two EFMs use different hydrogenases to reoxidise Fd(red). The figure shows how the requirement for non-zero DF for Conf_Hyd (**Eq. 1.1**) necessitates inefficient DF allocation to Hyd (**Eq. 1.2**) at pH_2_ = 10 kPa (red box). The range of value for Conf_Hyd correspond to a range of NADH/NAD^+^ values of 0.01 - 1 (-260 to -320 mV). Flux-weighting based on the stoichiometry arising from a 50:50 blend of Homoacetic and Equimolar. The flux-weighted DF (**A**) and FFE (**B**) of the two hydrogenases are shown as a function of Fd(red)/Fd(ox). A physiologically relevant range of NADH/NAD^+^ values is used to generate a range of solutions for Conf_Hyd ^34–36,38^. For illustration, a realistic range of Fd(red)/Fd(ox) are displayed, also in units of mV ^8,37.^

Taken together, it seems that blending EFMs to increase ATP yield at different pH_2_ is complicated by the fact that hydrogenases and other redox-balancing reactions, like RNF, cannot be combined without introducing substantial thermodynamic inefficiencies into a network.

## Discussion

Acidogenesis is typical of *Clostridial* glucose fermentation at neutral pH, with butyrate and acetate as major fermentation products ^3,5,6,39^. The specific molar ratio in which these acids are produced (Ace:But ratio) has been posited as a response to the pH_2_ in the fermenter’s environment (**Fig. 1**): increasing acetate production is accompanied by both increasing H_2_ yield and substrate-level phosphorylation (SLP). As SLP is the main source of ATP in anaerobic fermenters, it is thought that *Clostridia* flexibly adjust their product profile along a continuum of Ace:But ratios to maximise ATP production under the constraints of the pH_2_ in the environment: making more acetate, ATP, and H_2_ at lower pH_2_, and more butyrate (hence less H_2_) to maintain pathway DF at higher pH_2_ ^4^. This is not borne out by experimental observations, however, with fast-growing, glucose-limited anaerobic cultures showing strikingly stable Ace:But ratios despite changing pH_2_ (**Fig. S1**).

To explore why this might be the case, we first identified the three discrete product profiles which characterise acetate-butyrate production from glucose: Homoacetic (purely acetate production), Equimolar (equal parts acetate and butyrate), and Homobutyric fermentation (pure butyrate production). For the Equimolar and Homobutyric product profiles, multiple underlying stoichiometries were identified (**Fig. S2**). In order to identify these pathway stoichiometries, we decomposed a network of *Clostridial* glucose catabolism into elementary flux modes (EFMs). EFM enumeration is an unbiased way of identifying the “building blocks” of any redox-balanced metabolism ^31,40^.

*Clostridial* glucose catabolism can only reach product profiles intermediate to the three described above by “blending” EFMs. To produce more acetate than butyrate, a part of the glucose would have to pass via one EFM and a part via the other. We explored the thermodynamic efficiency of the EFMs and the blends using MDF analysis. For simplicity, only the most thermodynamically favourable (highest minimum DF) stoichiometry for each product profile was investigated in this study (**Fig. 2**).

MDF analyses attempt to identify ideal metabolite concentrations that maximise the thermodynamically limiting step – the step with the lowest DF ^29^. Flux-force efficacy (FFE), a value indicating how much of the total flux carried by an enzyme is being catalysed in the forward direction, scales hyperbolically as a function of DF. This has two implications: first, that reactions with low DF are catalysing lots of unproductive reverse flux, which can increase the amount of enzyme that needs to be made for that step to ensure adequate net forward flux; and second, that allocating very high (>10 kJ/mol) DF to a reaction does not substantially increase the efficiency of that step (FFE is virtually at its maximum at that point). In a sense, a MDF analysis is implicitly a resource allocation model, as it suggests that more resources (a higher enzyme cost) would have to be spent on reactions with low DFs, and that resource constraints would favour cells with more efficient DF allocation strategies ^41^. Noor *et al*. ^42^ have explicitly developed this idea, showing that despite the absence of detailed kinetic knowledge, pathway thermodynamics can give an indication of the enzyme levels in a cell.

We have also extended the MDF solver to include all feasible metabolite concentrations and DFs that yield the calculated minimal DF (**Fig. 3**). To do this, we calculated the MDF and then maximised and minimised all metabolite concentrations, reaction DFs, and redox ratios while keeping the MDF constant. This is based on how “flux variability analysis” extends “flux balance analysis” by first optimising the objective and then identifying the range of each variable that is possible for that solution ^43^.

Two of the EFMs identified in this study are known to occur experimentally – Homoacetic ^32,44^ and Equimolar ^3,8^. Although the Homobutyric stoichiometry had a reasonably favourable minimum DF and ATP yield, homobutyric glucose fermentation has not been reported in literature, as far as we could identify. While it has been suggested for the ruminal *Butyrivibrios* ^45^, this suggestion was based on genomic data and evidence of this product spectrum in practice was not presented. The reasons for this absence from the literature lie outside of the scope of our investigation and homobutyric fermenters might still be identified. *Clostridia* are widely accepted as butyrate producers, and butyrate is known to be important to gut and wider systemic health ^46^. The gut absorbs short-chain fatty acids like butyrate ^47^, which might abnegate some of the toxic effects of butyrate accumulation in that environment.

We have only investigated the effect of pH_2_ on the distribution of pathway DF. Other environmental variables, including the concentration of glucose, acetate, and butyrate in the environment will also have an effect *in situ*. Another key variable is temperature, which we have kept constant at 25°C in our analyses. For instance, changes in temperature could make Homoacetic feasible at higher pH_2_: the overall DF of this catabolic stoichiometry will increase 35 kJ per mole of glucose at 90°C (compared to standard conditions 25°C, 1M) ^8^.

### Stoichiometries yielding intermediate product profiles are inefficient

Our most striking result was that the Homoacetic+Equimolar blend of EFMs had a lower minimum driving force than either EFM at all pH_2_ (**Fig. 4**). The inefficiency held regardless of the proportion in which the EFMs are blended – combinations ranging from 5:95 to 95:5 all have lower minimum driving forces (DFs) than the EFMs (**Fig. 5**). No single enzyme was responsible for the result: two thirds of reactions had FFEs approximately 2-fold lower than both EFMs (**Fig. S3**). Using literature-derived k_cat_ and molecular weights, this can be reinterpreted as enzyme cost (**Fig. 7**). This indeed reflects a significant increase in enzyme cost for the Homoacetic+Equimolar blend, compared to either Homoacetic or Equimolar fermentation, underscoring the trade-off between ATP yield and enzyme cost.

**Figure 7.**
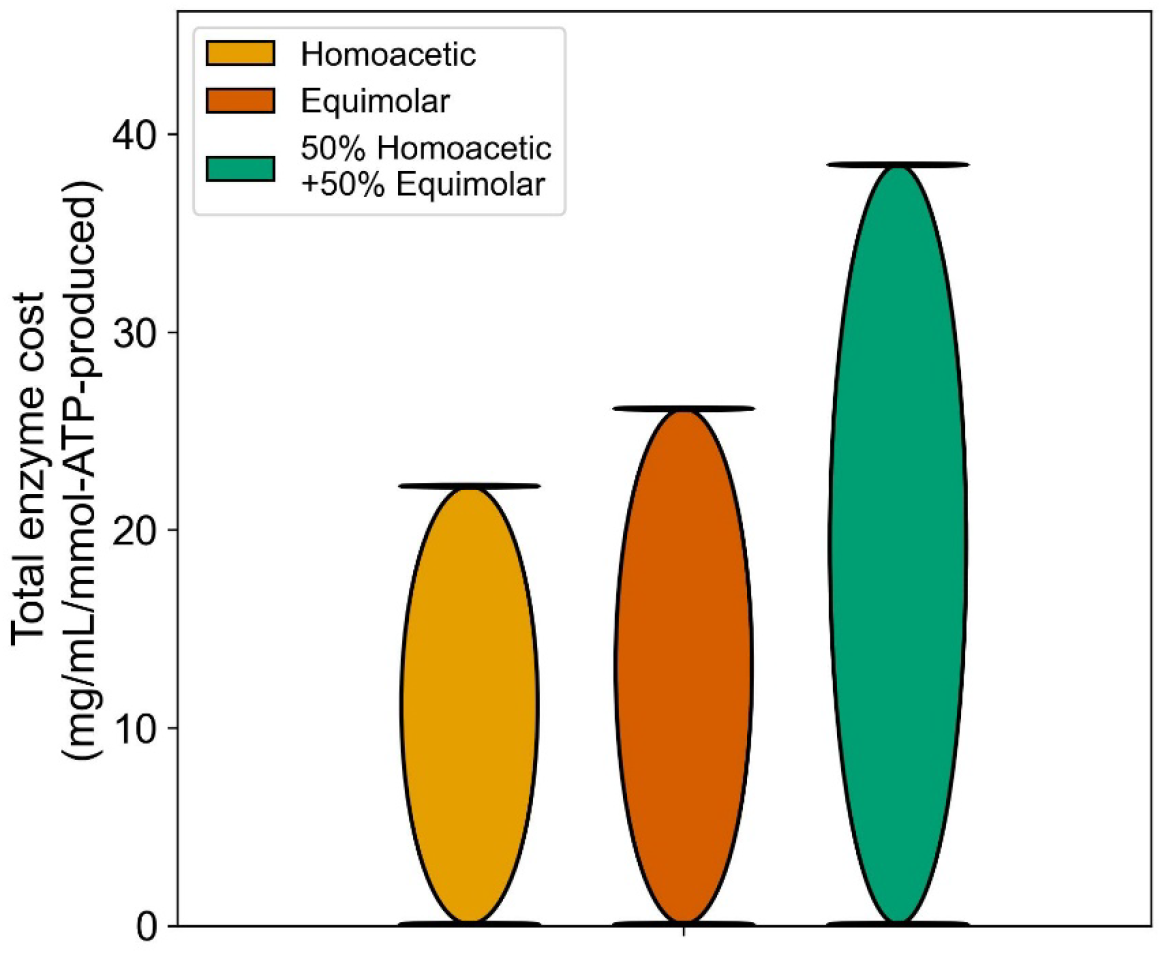
Enzyme cost per ATP produced based on reaction driving forces. Using literature-derived *Clostridial* k_cats_ and molecular weights, the pathway DFs can be reinterpreted in terms of hypothetical enzyme costs (**Table S6**). This assumes full kinetic saturation and no allosteric inhibition (see *Methods* for calculation). This example pertains to the pathway DFs at pH_2_ = 10 kPa.

This increased enzyme cost of blended EFM stems from the need to run both Hyd and Conf_Hyd in the forward direction. Due to them sharing Fd(red)/Fd(ox) as redox pair, Conf_Hyd cannot run on the forward direction without increasing the Fd(red)/Fd(ox) ratio to well above 10 (around –440 mV; **Fig. 6**). This forces more than 20 kJ/mol of driving force to be allocated to Hyd, which decreases the total DF available for other reactions **Box 1**. The knock-on effects of this go beyond just the hydrogenases. Other enzymes (PFOR and BCD-ETF) also have Fd(red) and Fd(ox) as product-substrate pair, meaning that their DF would be very unfavourably affected by such a high ratio. Other metabolites might increase/decrease to compensate, but this is limited by the concentration constraints imposed on the MDF solver. We have developed this hypothesis for the case Homoacetic+Equimolar, the two catabolic stoichiometries known to occur in literature (**Fig. 5**).

### Ace:But ratios in nature

Our results provide a possible explanation for the observed insensitivity of the Ace:But ratio in fast-growing, glucose-limited anaerobes to changes in pH_2_ (**Fig. S1**). One might speculate that cultures growing at lower dilution rates need to make fewer ribosomes to divide and grow fast, leaving more space on the proteome to compensate for thermodynamically inefficient, but higher ATP-yielding, catabolisms ^48^. This might explain the increased responsiveness to pH_2_ of these slower-growing cultures. At higher dilution rates, however, the enzyme cost incurred by these inefficiencies might prevent microbes from growing fast enough to avoid wash-out.

Experimentally, Ace:But ratios intermediate to the product profiles of the EFMs are encountered, but probably stem from the interaction of catabolism and anabolism ^35,39,49–51^. A product profile is not only a result of ATP-producing glucose catabolism. Rather, it is a mixture of complex interactions between anabolism and catabolism where not only ATP but also NADH (for example as an anabolic byproduct of amino acid synthesis via the oxidative phase of the TCA cycle ^2^) and NADPH (an important electron donor to, for example, fatty acid synthesis ^52^) are exchanged. The transfer of electrons to or from the biomass has been suggested to be a major determinant of fermentation product profiles^53^. What is more, microbes in the lab are often not growing on glucose alone, but on media containing undefined components like yeast extract, and do not produce acetate and butyrate only.

## Conclusions

We have performed an MDF analysis suggesting that stoichiometries that combine specific redox-balancing reactions decrease their thermodynamic efficiency. This analysis has led to our hypothesis that *Clostridial* glucose fermentation at neutral pH might not be a continuum of fractional combinations of acetate and butyrate, but a discrete set of thermodynamically optimal stoichiometries. This implies that changing pH_2_ does not necessarily translate to a change in product profile. This is relevant for anaerobic cultivations in which a given product must be maximised, e.g. butyrate or H_2_ gas. It is also of consequence for synthetic biology, implying that enzymes cannot be recombined purely based on pathway yield, but must also be thermodynamically compatible.

## Methods

### Construction of Clostridial reaction network

Following on the simplified network posited by Spormann^4^ we constructed a generic network of all reactions involved in the pH-neutral acidogenic fermentation from glucose, and limited to acetate, butyrate, H_2_, and CO_2_ as products. The network included 34 reactions – 10 exchanges and 24 conversions – and considered only one cytosolic compartment. Reaction reversibility was derived from the UniProt database ^54^. Metabolites ATP, ADP, Pi, and the considered translocated protons across the cellular membrane (H^+^) were not internally balanced, as these can be exchanged with other metabolic pathways *in vivo*.

Some *Clostridia* make use of a phosphotransferase systems (PTS) to simultaneously import and phosphorylate glucose ^2^, while others have separate import and phosphorylation steps, the latter catalysed by a hexokinase (HK) enzyme ^55^. Our generic network contained two separate steps.

In addition to the redox-balancing reactions presented by Spormann^4^ – the confurcating hydrogenase (Conf_Hyd), hydrogenase (Hyd), and RNF – we also include the energy-converting hydrogenase (EC_Hyd) ^9^. Since the RNF and EC_Hyd can both produce a proton-motive force, we also include literature-derived PMF-producing stoichiometries for those reactions as well as an ATP synthase (ATPase) ^8,11^.

The full network is available as supplementary **Table S2**.

### Enumeration of elementary flux modes

Elementary flux mode (EFM) enumeration was performed on the network in **Table S2**. Then, the EFMs were normalised to glucose consumption, and are presented in **Table S3**.

### Thermodynamic data

Standard Gibbs free energies (ΔG°’) were obtained using the eQuilibrator API ^56,57^ at pH = 7.4, pMg = 3, and ionic strength = 0.1 M. A confidence interval of 99% was chosen.

### H_2_ gas-liquid transfer

H_2_ is assumed to partition between the gas and liquid phases according to ideal equilibrium behavior, governed by Henry’s law with a constant Henry coefficient, so the pH_2_ indicated in the figures pertains directly to the H_2_ concentration in the liquid phase.

### Max-Min driving force

We applied <u>M</u>ax-Min <u>D</u>riving <u>F</u>orce (MDF) analysis to determine the feasibility of the catabolic EFMs and combinations of EFMs. This was done according to the method of Noor *et al*. ^29^, formally defined as:

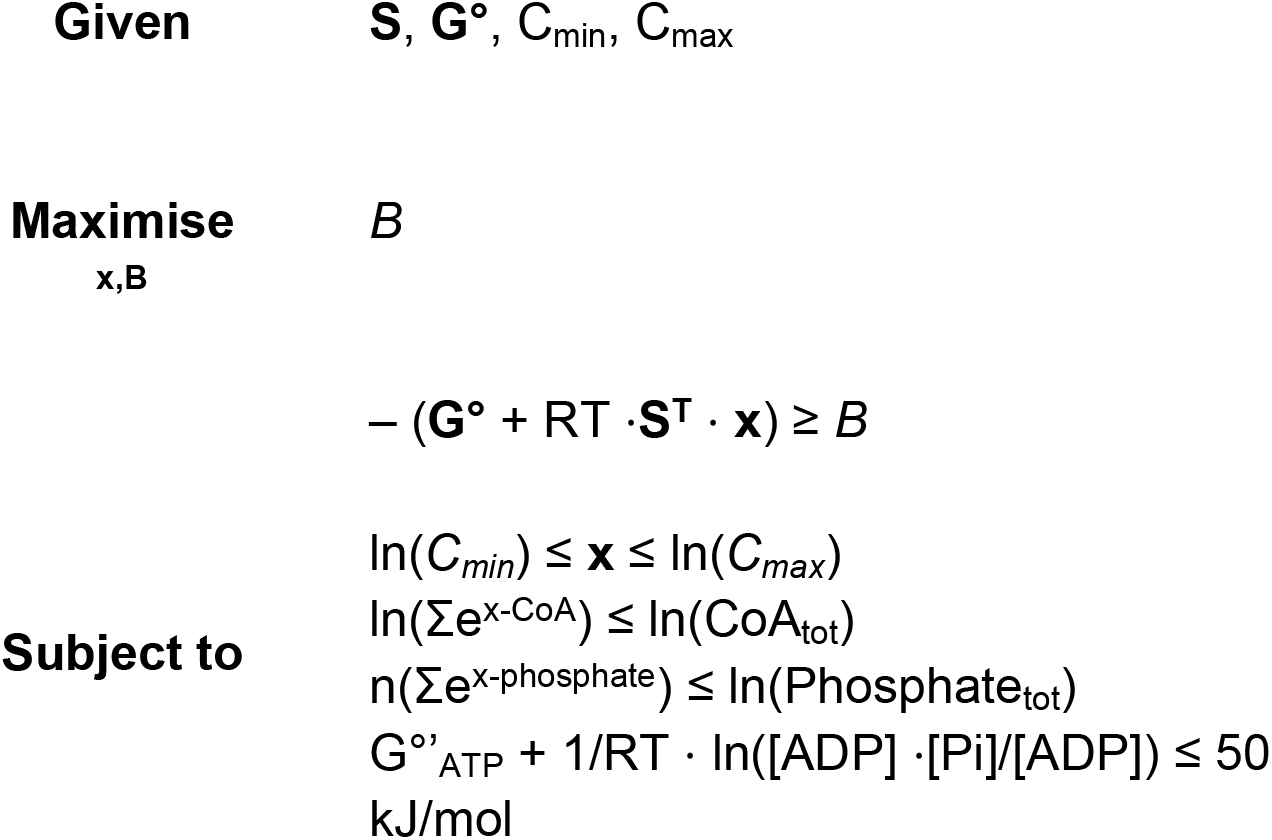

Where *B* is a tight lower bound – the minimum – on the driving forces of all pathway reactions. The vector of the driving forces is given by *– (****G°*** *+ RT ⋅* ***S***^***T***^ *⋅* ***x***). Within that term, ***S*** is a stroichiometric matrix of shape *i×j* containing *i* metabolites and *j* reactions, ***G°*** is a column vector containing standard Gibbs free energy changes for *j* reactions as obrained from eQuilibrator, and **x** is a column vector of the natural logarithms of the *i* metabolite concentrations in the model. *C*_*min*_ and *C*_*max*_ are concentration bounds for model metabolites and were set to 1 µM and 10 mM, respectively. The only exceptions were H_2_O, which was set to a constant concentration of 1 M, free P_i_, which had an upper bound equal to the total phosphate pool size (50 mM) and free CoA, which had an upper bound equal to the total CoA pool size (1.5 mM). These bounds are found in **Table S4**. External metabolite concentrations (Gluc_out, Ace_out, But_out and CO2_out) were all set 1 mM. Only H_2_ was set to the defined and explicitly indicated values.

The ratios of Fd(red)/Fd(ox) and NADH/NAD^+^ were not fixed unless explicitly indicated, but bounded like all other metabolites considered.

Two moiety conservation constraints were also imposed on the optimisation: co-enzyme A (CoA) and phosphate conservation. To maintain the constraint in a convex form, moiety conservation was implemented as an inequality constraint as presented in González-Cabaleiro *et al*.^58^: The sum of moiety-containing metabolites was not allowed to exceed a set upper scalar bound, 1.5 mM for CoA ^59^ and 50 mM for phosphate ^60^.

Practically, the following constraints were applied:

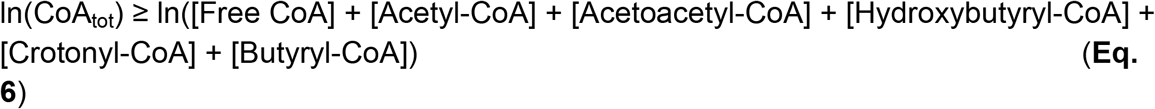

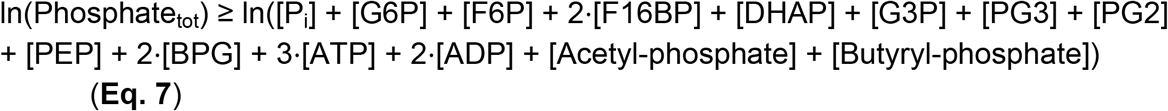

A total adenylate pool size of 5 mM was chosen and a constant ratio of ATP/ADP set to 10/1, typical of growing cells ^10^. Effectively, the ATP concentration was set to ca.

4.54 mM and the ADP concentration to ca. 0.454 mM. Δ_r_G of ATP hydrolysis is function of the variable P_i_ concentration, therefore, it is also a variable for which we impose an upper bound of –50 kJ/mol ^61^ (or +50 kJ/mol for ATP formation). Proton-motive force (PMF) was not an explicit variable in the MDF stoichiometries. To represent the energetic cost of proton-pumping by EC_Hyd (1 proton per reaction, equivalent to 0.25 ATP) and RNF (2 proton per reaction, equivalent to 0.5 ATP) ^8,11^, the Δ_r_G of ATP formation was added to the Δ_r_G calculations of those reactions.^4^

The parameters of the optimisation are contained in **Table S5**.

### Variability within MDF solution

After determining the minimum DF, we carried out a three-step variability analysis to identify all metabolite concentrations, redox ratios, and reaction DFs for which the MDF solution holds. In the first step, the calculated *B* was set as a lower bound on all pathway DFs. Then, each metabolite concentration was maximised and minimised

**Step 1: Concentration variability:**

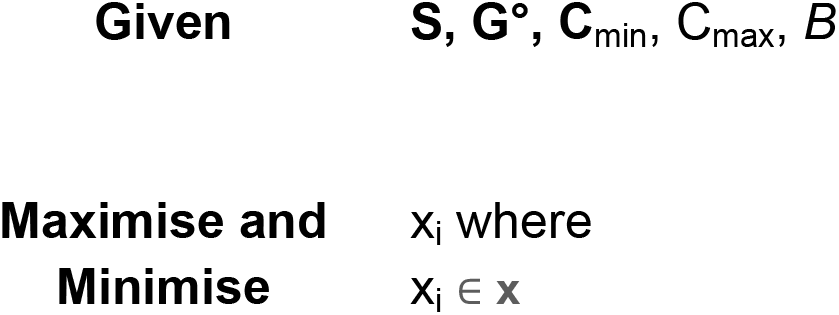
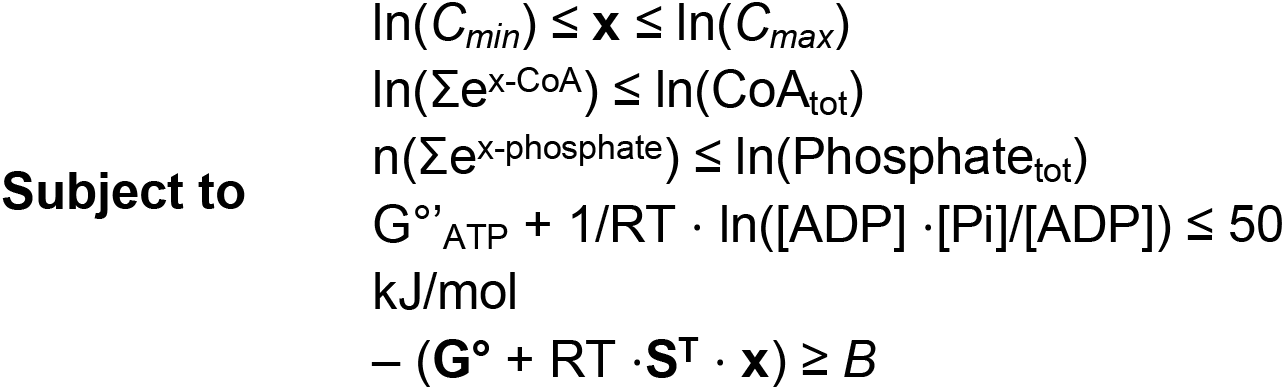

**Step 2: Redox ratio variability**

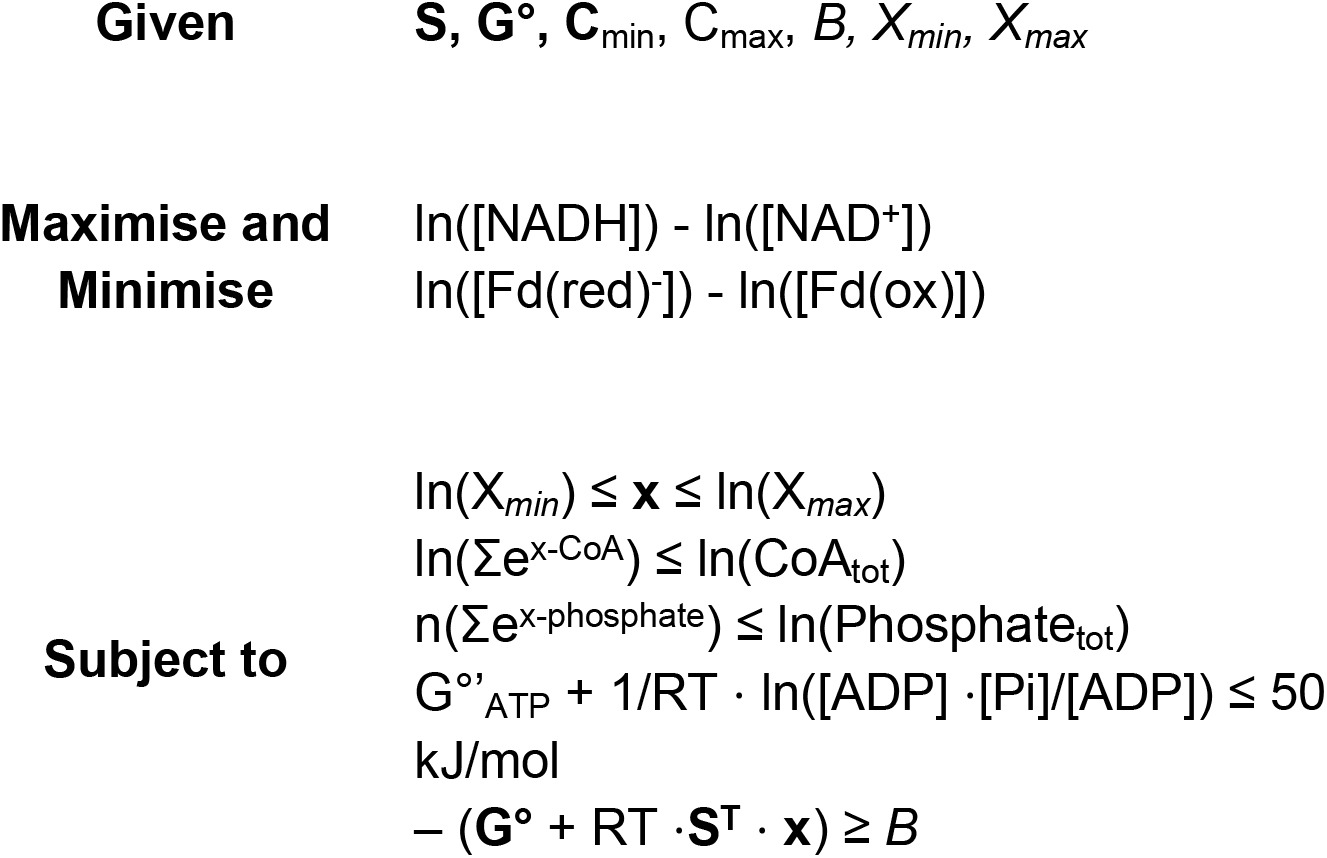

Where **X**_**min**_ and **X**_**max**_ are column vectors containing the maximum and minimum concentration of each pathway metabolite.

**Step 3: Driving force variability**

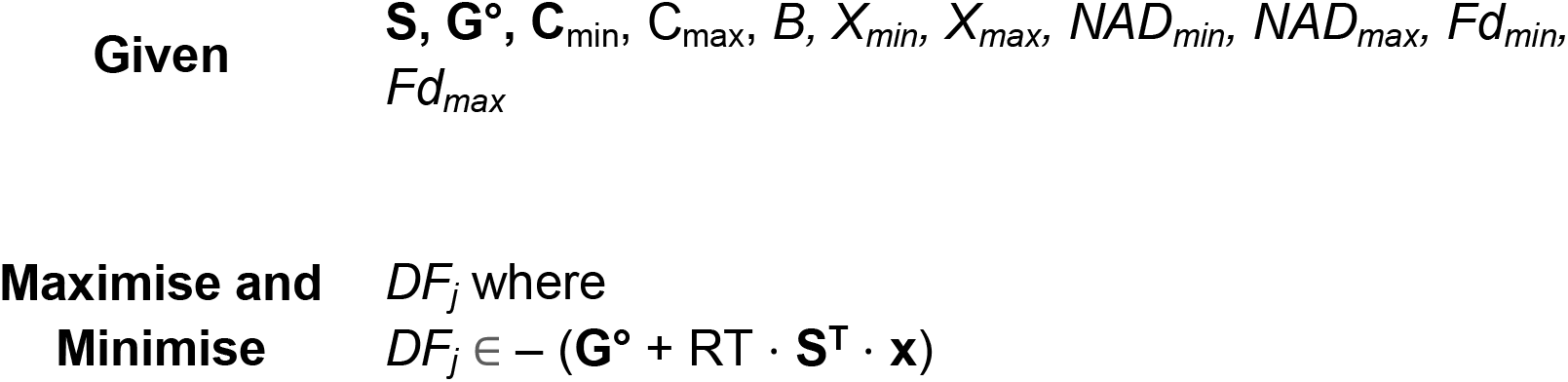
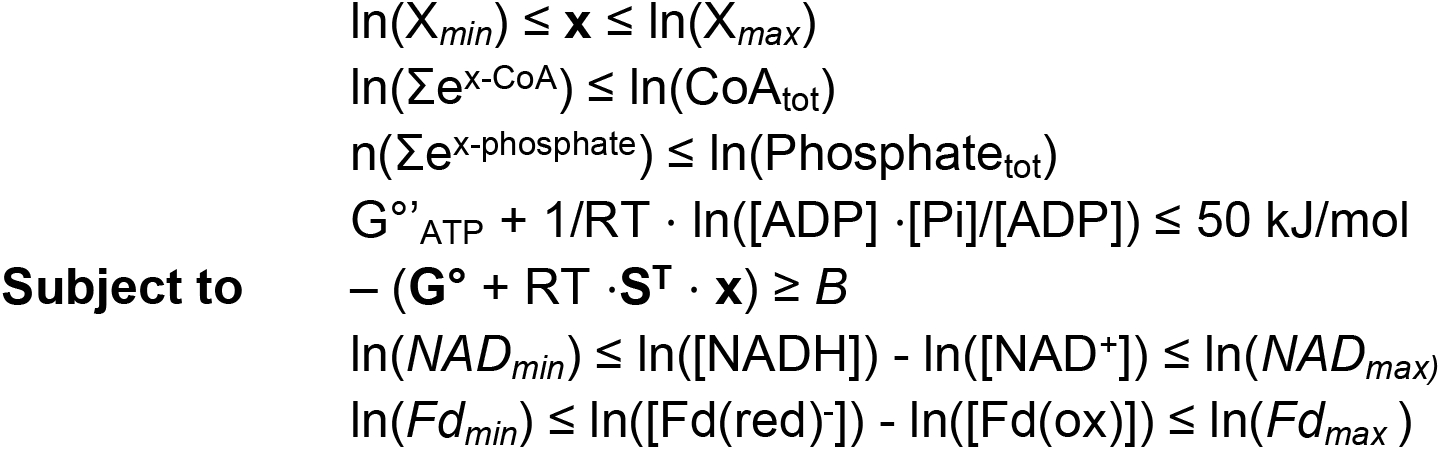

Where *NAD*_*min*_ and *NAD*_*max*_ are the min- and maximum ratios of NADH/NAD^+^, respectively, and *Fd*_*min*_ and *Fd*_*max*_ for Fd(red)/Fd(ox). *DF*_j_ representes the driving force of one reaction within the colum vector of driving forces calculated by: – (**G°** + RT ⋅ **S**^**T**^ ⋅ **x**).

### Enzyme cost

Total catabolic enzyme demand was calculated using the formula from reversibility-based enzyme cost minimisation (ECM2) as proposed by Noor *et al*. ^42^:

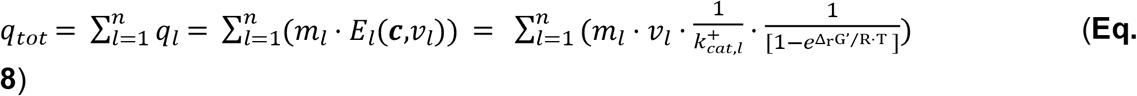

where:

- *q*_*tot*_ is the total pathway enzyme cost (g/L)
- *q*_*l*_ is the enzyme cost of one reaction (g/L)
- *m*_*l*_ is the mass of the enzyme (Da)
- *E*_*l*_ is the enzyme level of one reaction (M)
- ***c*** is the metabolite concentrations (M)
- *v*_*l*_ is the rate of reaction l (mol/s/g-enzyme)
- *k*^*+*^_*cat*_ is the catalytic constant of reaction l (s^-1^)
- Δ_r_G’/RT

Note that *k*^*+*^_*cat*_ is usually expressed per active site. We multiply it by the number of active sites.

We set *v*_*l*_ so that the fluxes represent the steady state that would yield 1 mmol of ATP. This is done by dividing the flux through each reaction by the mmol yield of ATP on glucose (1000*Y^ATP^_Gluc_). This means that the *q*_*tot*_ (and all the constituent q_l_) can be expressed in units of g/L/mmol-ATP-produced or mg/mL/mmol-ATP-produced.

ECM2 effectively assumes that the enzymes are optimally substrate-saturated (*η*^*sat*^_*l*_(***c****) =1*) and that regulation is optimally forward (*η*^*reg*^ _*l*_ (***c****) = 1*) and calculates enzyme cost only based on k_cat_ and the displacement of the reaction from equilibrium.

We did not minimise enzyme cost, but rather calculated the optimal driving force distribution with only concentration constraints and then calculated the hypothetical enzyme cost associated with it as an illustration. The kinetics and results of these calculations are shown in **Table S6**.

### Software

**Table S2** was converted from an XLSM file to an XML file and constraint-based checks run for elemental and charge balance using COBRApy v. 0.3.1 ^62^. EFMs were enumerated using the python package efmtool v. 0.2.1 ^30^. Convex optimisation was carried out using CVXPY.

## Data availability statement

Data, scripts, and notebooks used to execute the model and generate the paper figures are stored and made available on GitHub (https://gitlab.tudelft.nl/codendaal/thermodynamic-pathway-model/-/tree/3ba399ec8ce3b018775301732bd8c20fdf725dec/).

## Acknowledgements

The authors would like to thank Dr Robbert Kleerebezem for many thoughtful discussions and valuable insights. The figures were compiled in with BioRender.com. This work was supported by the European Union’s Horizon research and innovation programme under Grant Agreement No 10113540 (<u>Nutritive</u>), which was awarded to Dr

R. González-Cabaleiro.

## Supplementary Figures

**Figure S1.**
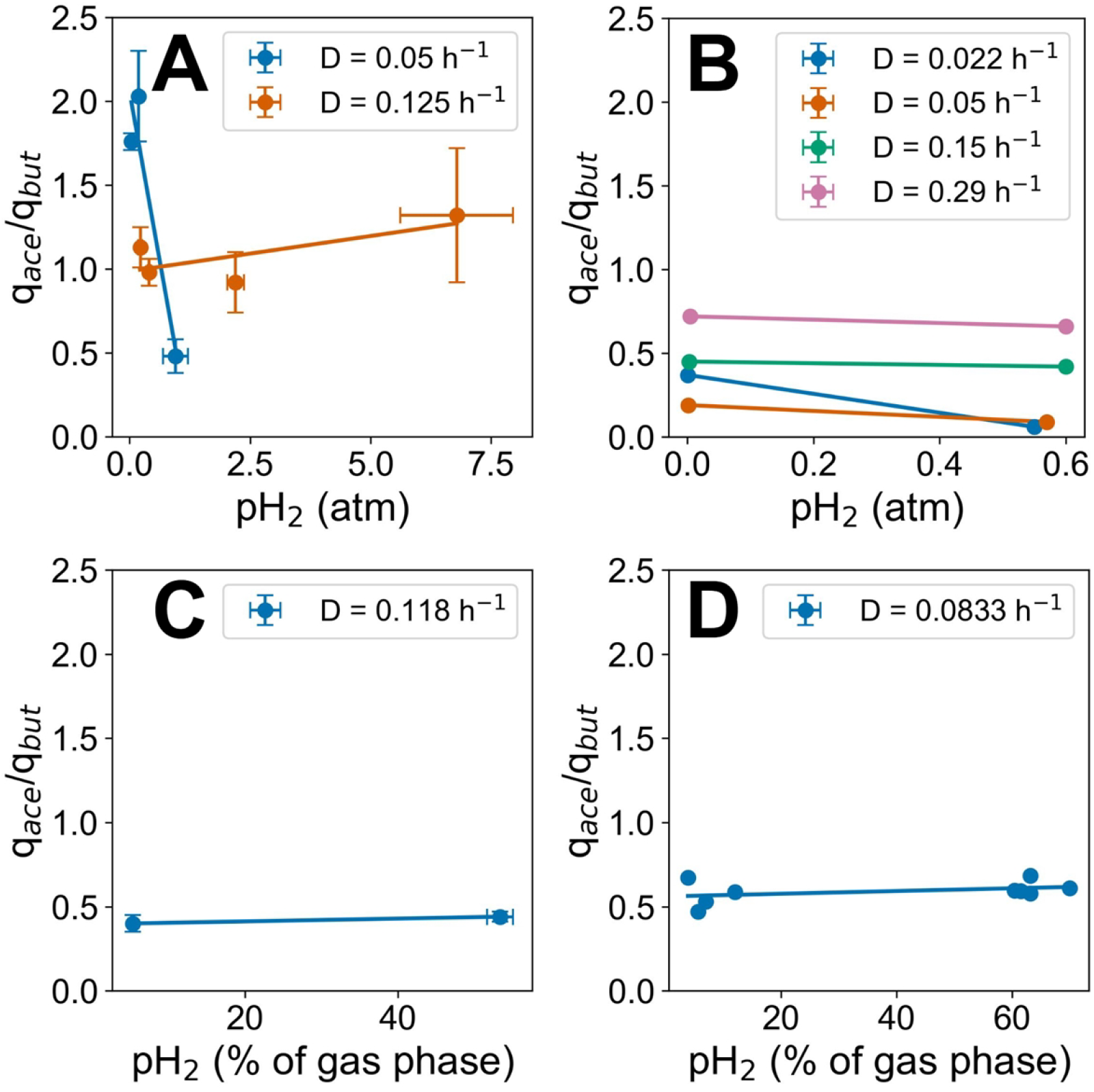
The ratio q_ace_/q_but_ as a function of pH_2_ in literature. Acetate *versus* butyrate production ratio of glucose limited cultures as a function of pH_2_ in the units used in the original publications. The cultures were described as mixed ^24^ (**A**), pure *Clostridium butyricum* ^26^ (**B**), hydrogen-producing mixed ^25^ (**C**), and wastewater sludge grown on a sucrose medium ^27^ (**D**). Data in (**D**) is presented as individual data points because they were experiments run at different gas flow rates. Dilution rates Details available in **Table S1**.

**Figure S2.**
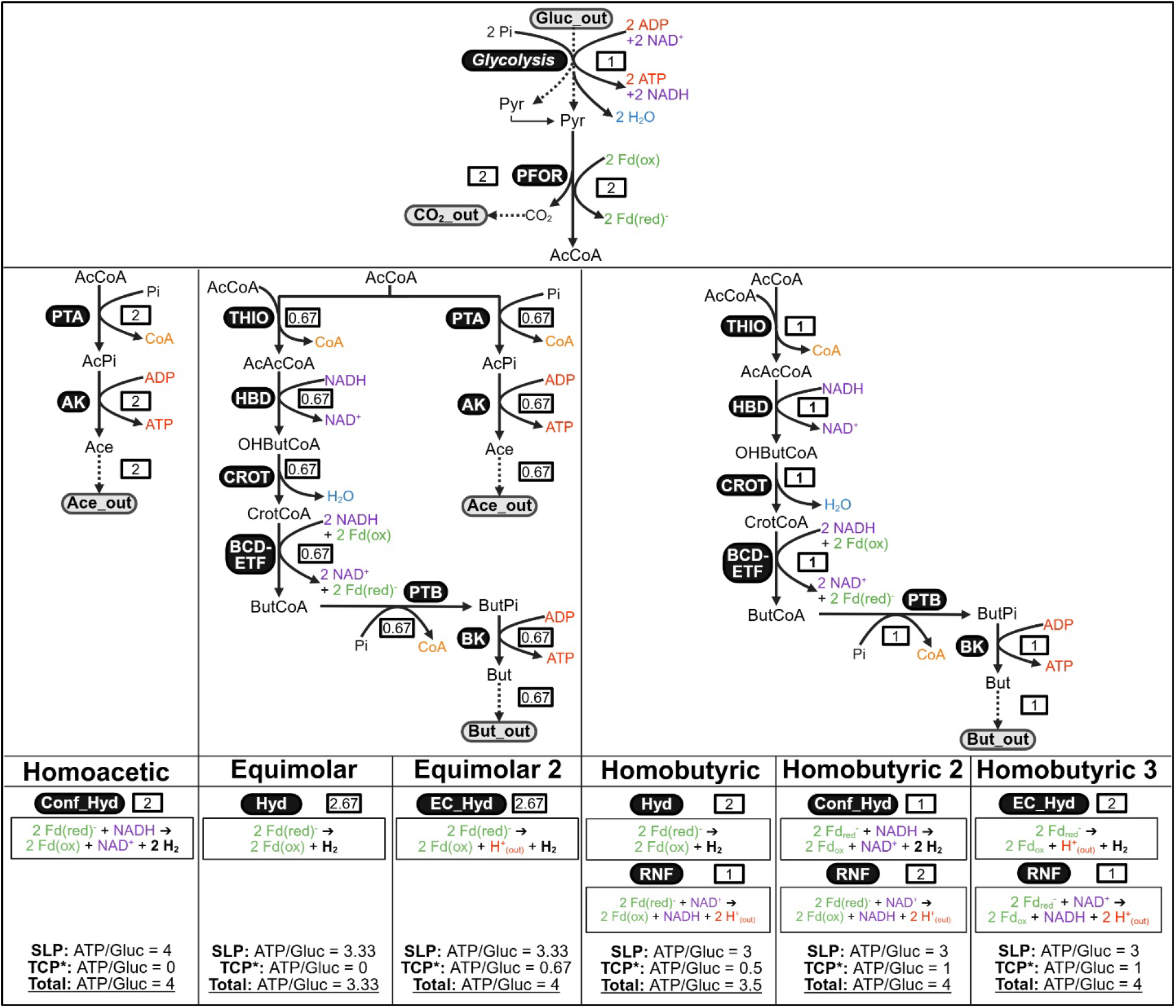
All elementary flux modes in *Clostridial* acidogenesis. The decomposition of the network reactions typically associated with *Clostridial* acidogenesis yielded six elementary flux modes (EFMs). They are displayed relative to the flux of glucose consumption (white boxes indicate the relative flux of each reaction). All of these EFMs contained the ten reactions of glycolysis plus pyruvate:ferredoxin oxidoreductase (PFOR), indicated at the top of the figure. This is indicated as a dashed line with its conversion within each EFM, again relative to glucose consumption flux. The six EFMs are displayed vertically. They are named according to their acid products: pure acetic acid production (Homoacetic), equimolar production of acetic and butyric acid (Equimolar), and pure butyric acid production (Homobutyric). ATP and H_2_ production can vary based on which redox-balancing reactions are active, yielding the variants of equimolar and homobutyric acidogenesis that are distinguished by the six panels at the bottom of the figure. Black, curved boxes contain enzyme name abbreviations as defined in the glossary and grey curved boxes

**Figure S3.**
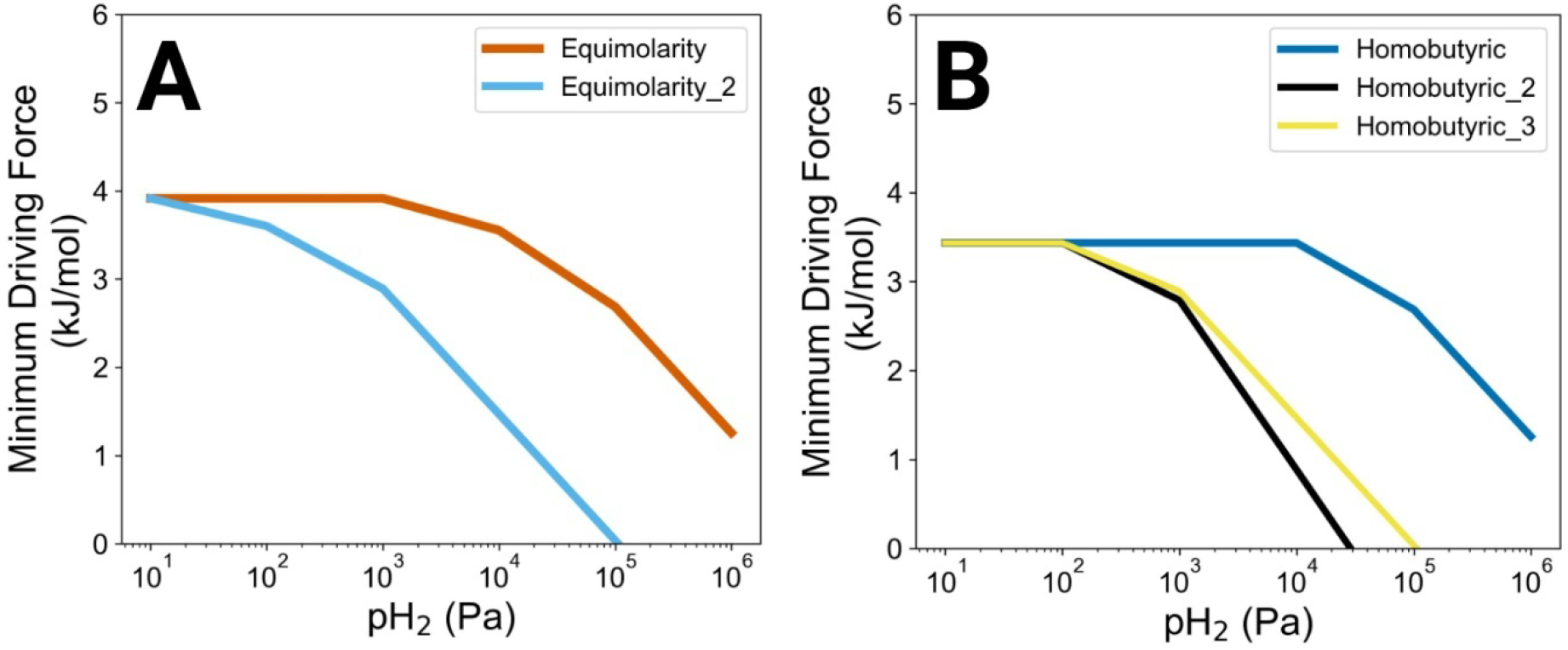
Minimum driving force as a function of pH_2_ in stoichiometries associated with Equimolar and Homobutyric. Lowest DF (kJ/mol) of any reaction stoichiometry, the thermodynamic bottleneck of the pathway. Result of MDF. **A**. Equimolar stoichiometries. **B**. Homobutyric stoichiometries.

**Figure S4.**
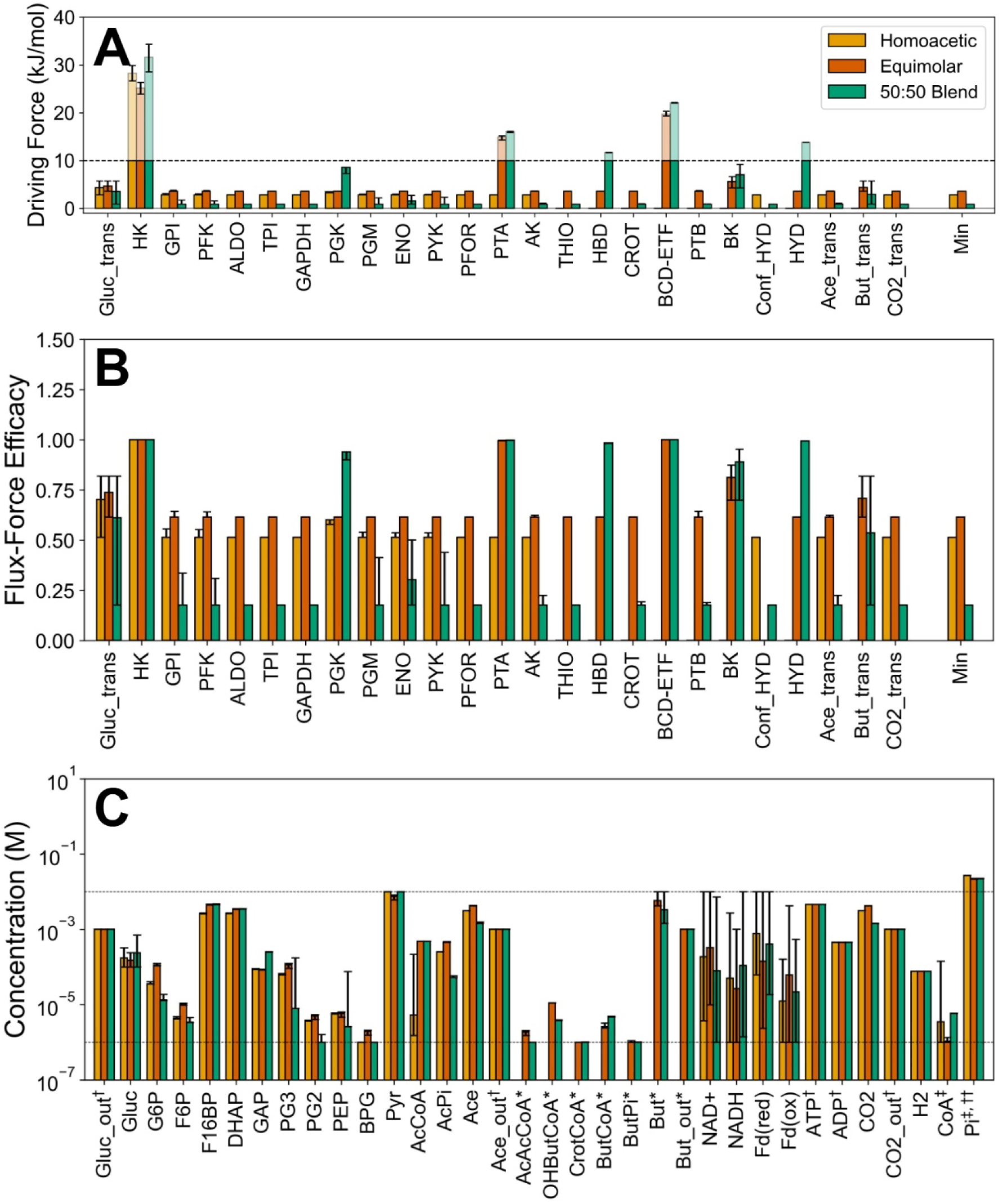
Effect of blending Homoacetic and Equimolar per reaction and per metabolite. MDF result for Homoacetic, Equimolar, and a 50:50 combination (pH_2_ = 10 kPa). Single asterisks (*) indicate where the reaction was not present in Homoacetic while double asterisks (**) indicate where it was only present in Equimolar. **A**. Driving force (DF) allocated to each reaction. **B**. The flux-force efficacy (FFE) calculating from the driving forces in panel. **C**. Metabolite concentrations. ^†^ refers to fixed metabolite concentrations, ^††^ to metabolites that are not constraint to concentrations between 1 µM and 10 mM, and ^‡^ to the free version of a conserved moiety.

**Figure S5.**
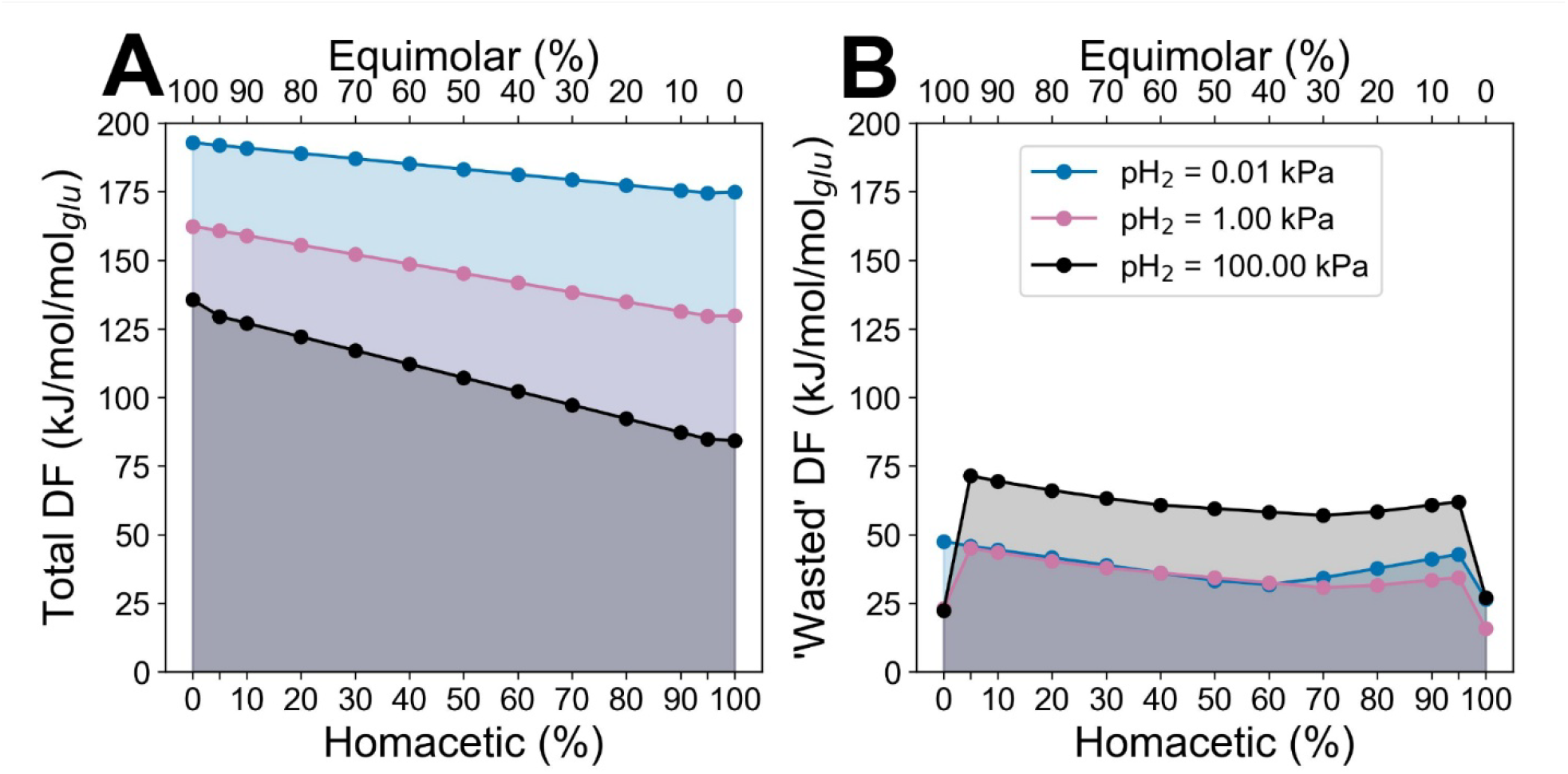
Effect on driving forces of blending Homoacetic and Equimolar. Homoacetic and Equimolar were blended in different ratios and an MDF analysis performed at three different pH_2_ value. The flux-weighted total (**A**) and ‘wasted’ DF (**B**) are shown. Note the two x-axes representing the % of Equimolar (top) and Homoacetic (bottom).

**Figure S6.**
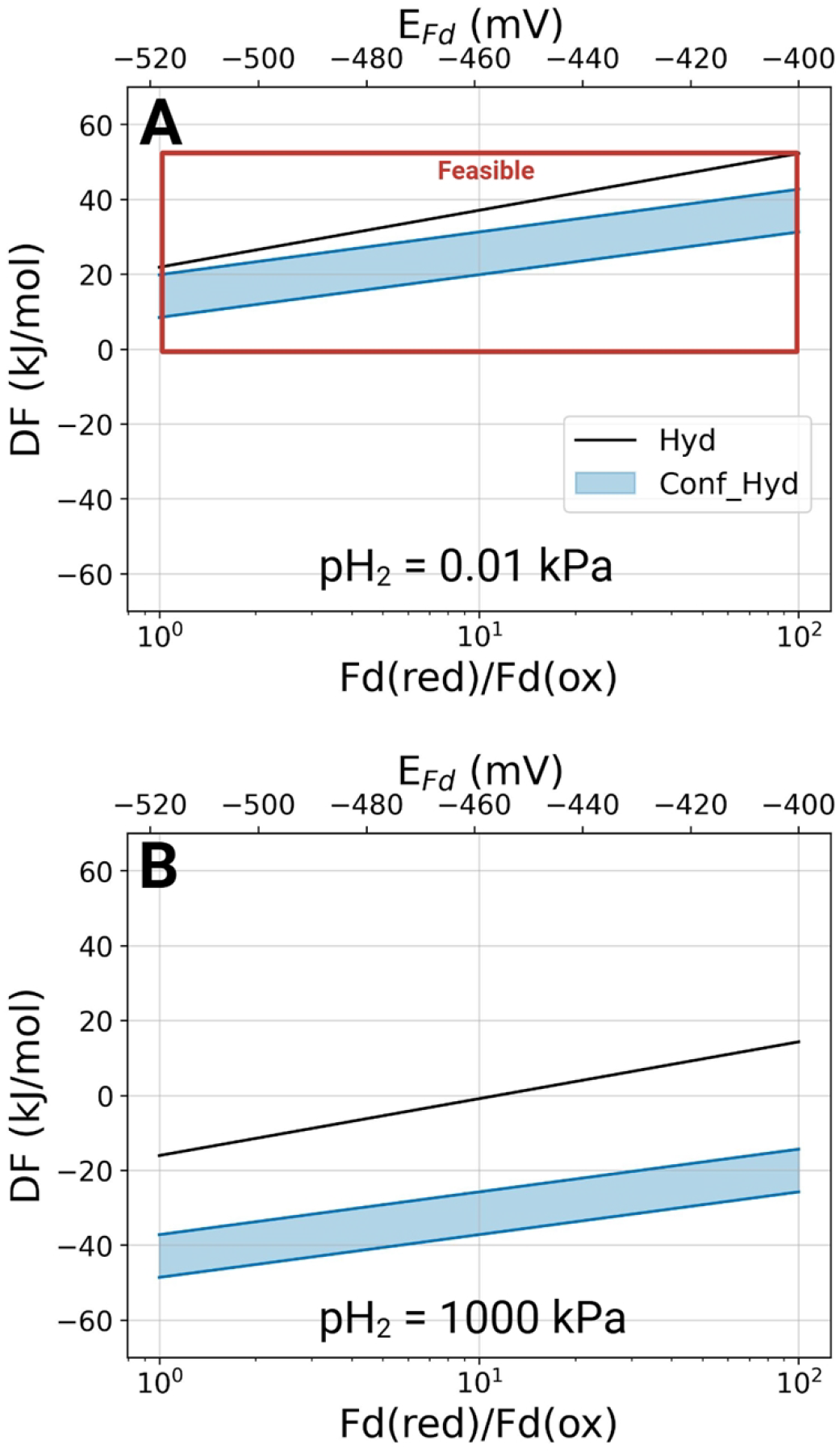
Combining Hyd and Conf_Hyd introduces thermodynamic limitations at different pH_2_. The figure shows the correlation between the flux-weighted DF of Hyd and Conf_Hyd at pH_2_ = 0.01 kPA (**A**) and pH_2_ = 1000 kPa (**B**). Flux-weighting based on the stoichiometry arising from a 50:50 blend of Homoacetic and Equimolar_II. A realistic range of NADH/NAD^+^ and of Fd(red)/Fd(ox) are tested, also in units of mV.

**Figure S7.**
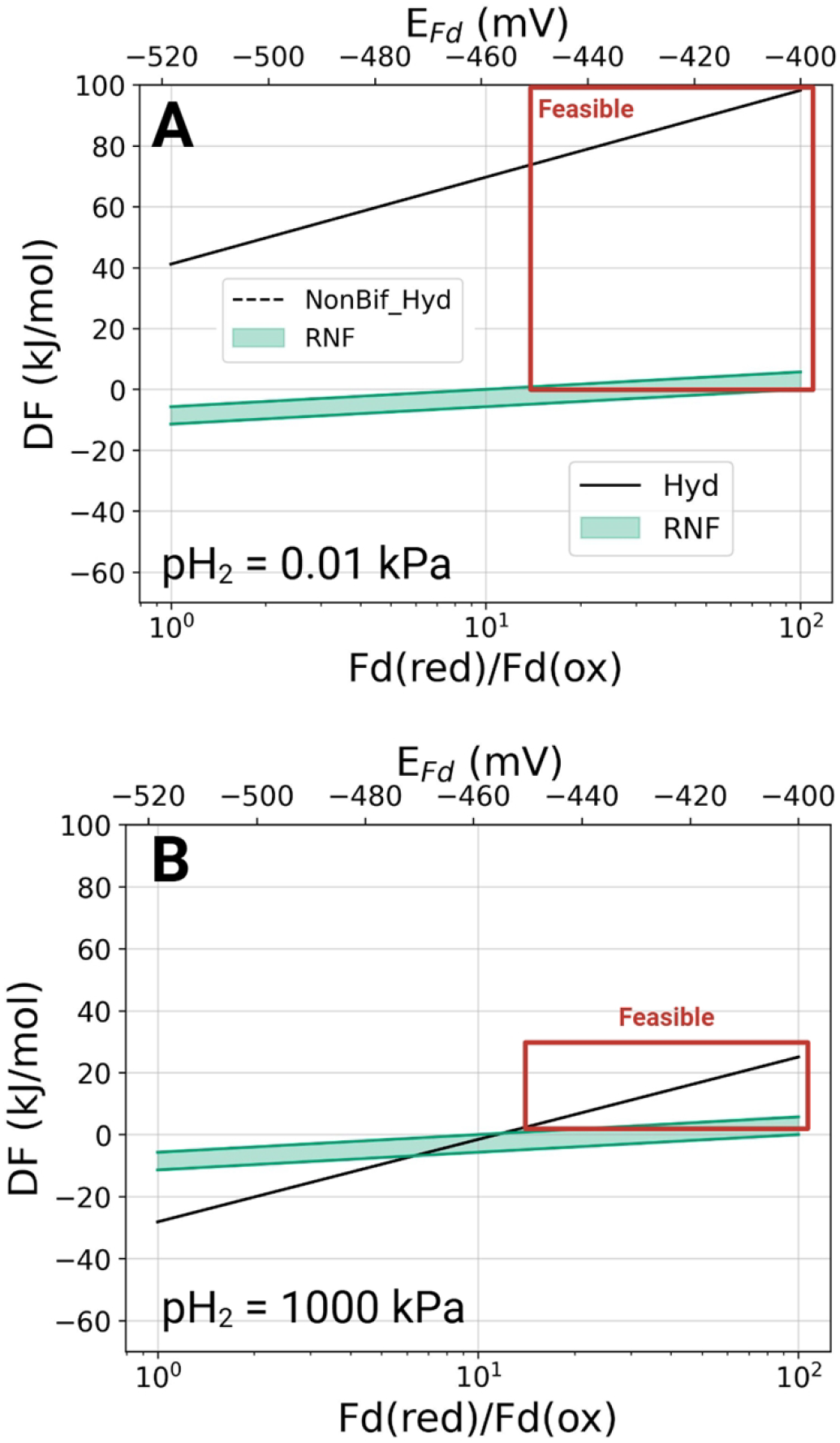
Combining Hyd and RNF introduces thermodynamic limitations at different pH_2_. The figure shows the correlation between the flux-weighted DF of Hyd and RNF at pH_2_ = 0.01 kPA (**A**) and pH_2_ = 1000 kPa (**B**). Flux-weighting based on the stoichiometry arising from a 50:50 blend of Homobutyric and Equimolar. A realistic range of NADH/NAD^+^ and of Fd(red)/Fd(ox) are tested, also in units of mV.

## Supplementary Tables

Table S1 - Ace-But ratio in literature.xlsx

Table S2 - EFM Stoichiometries.xlsx

Table S3 - Costridial catabolism.xlsx

Table S4 - Metabolites and concentration bounds.xlsx

Table S5 - MDF parameters.xlsx

Table S6 - Kinetics for enzyme cost calculation.xlsx

